# Waste not, want not: Revisiting the analysis that called into question the practice of rarefaction

**DOI:** 10.1101/2023.06.23.546312

**Authors:** Patrick D. Schloss

**Author notes:** To whom correspondence should be addressed Department of Microbiology & Immunology University of Michigan Ann Arbor, MI 48109.

## Abstract

In 2014, McMurdie and Holmes published the provocatively titled, “Waste not, want not: why rarefying microbiome data is inadmissible”. The claims of their study have significantly altered how microbiome researchers control for the unavoidable uneven sequencing depths that are inherent in modern 16S rRNA gene sequencing. Confusion over the distinction between the definitions of rarefying and rarefaction continues to cloud the interpretation of their results. More importantly, the authors made a variety of choices in designing and analyzing their simulations that were potentially problematic. I identified 11 factors that could have compromised the results of the original study. I reproduced the original simulation results and assessed the impact of those 11 factors on the underlying conclusion that rarefying data is inadmissible. Throughout, the design of the original study made choices that caused rarefying and rarefaction to appear to perform worse than they truly did. Most important were the approaches used to assess ecological distances, the removal of samples with low sequencing depth, and not accounting for conditions where sequencing effort is confounded with treatment group. Although the original study criticized rarefying for the arbitrary removal of valid data, repeatedly rarefying data many times (i.e., rarefaction) incorporates all the data. In contrast, it is the removal of rare taxa that would appear to remove valid data. Overall, I show that rarefaction is the most robust approach to control for uneven sequencing effort when considered across a variety of alpha and beta diversity metrics.

**Importance:** Over the past 10 years, the best method for normalizing the sequencing depth of samples characterized by 16S rRNA gene sequencing has been contentious. An often cited paper by McMurdie and Holmes forcefully argued that rarefying the number of sequence counts was “inadmissible” and should not be employed. However, I identified a number of problems with the design of their simulations and analysis that could compromise their results. In fact, when I reproduced and expanded upon their analysis, it was clear that rarefaction was actually the most robust approach for controlling for uneven sequencing effort across samples. Rarefaction limits the rate of falsely detecting and rejecting differences between treatment groups. Far from being “inadmissible”, rarefaction is a valuable tool for analyzing microbiome sequence data.

## Introduction

Microbiome studies that use amplicon sequencing to characterize multiple samples use PCR to amplify 16S rRNA gene fragments using primers with distinct barcodes or index sequences (1–3). These barcodes allow researchers to pool PCR products and then deconvolute the resulting sequence data based on the barcode sequences. Despite researchers’ best efforts to generate equimolar pools of PCR products, it is common to observe more than 10-fold variation in the number of sequences per sample (4). Researchers desire strategies to minimize uneven sequencing depth and thus need methods to control for this unevenness in their analyses. Of course, uneven sequencing is not unique to microbiome research and is a challenge faced by all community ecologists (5, 6). Common approaches to control for the effects of uneven sequencing depths include use of proportional abundance (i.e., relative abundance), scaling of counts, parameter estimation, and rarefaction (7–23).

In 2014 Paul McMurdie and Susan Holmes published “Waste not, want not: why rarefying microbiome data is inadmissible” (WNWN) in *PLOS Computational Biology* (24). Their paper has had a significant impact on the approaches that microbiome researchers use to analyze 16S rRNA gene sequence data. According to Google Scholar, this paper has been cited more than 2,400 times as of June 2023. There has been a rebuttal of WNWN that showed how rarefaction is beneficial in some cases (25); however, the proponents of WNWN appear to be holding sway over the microbiome community. I have received correspondence from researchers over the past 10 years asking how to address critiques from reviewers who criticize my correspondents’ analysis for using rarefaction (e.g., see https://twitter.com/inanna_nalytica/status/1264299305067786242). I have also received similar comments from reviewers regarding my own work. For example, I received the critique for a preprint that I posted in 2020 that analyzed the practice of removing rare taxa from microbiome analyses [see comments linked to (4); https://web.archive.org/web/20230308191505/http://dalmug.org/filter-review/]. In the process of responding to these reviewers’ comments and preparing a manuscript investigating rarefaction and other approaches to control for uneven sequencing effort, I grew to appreciate that there is significant confusion in the field over what is meant by “rarefying” and “rarefaction”.

McMurdie and Holmes discussed and defined “rarefying” in WNWN (citation numbers updated from the original to those of the current study):

> “Instead, microbiome analysis workflows often begin with an ad hoc library size normalization by random subsampling without replacement, or so-called rarefying (13, 26, 27). There is confusion in the literature regarding terminology, and sometimes this normalization approach is conflated with a non-parametric resampling technique — called rarefaction (5), or individual-based taxon re-sampling curves (28) — that can be justified for coverage analysis or species richness estimation in some settings (28), though in other settings it can perform worse than parametric methods (29). Here we emphasize the distinction between taxon re-sampling curves and normalization by strictly adhering to the terms rarefying or rarefied counts when referring to the normalization procedure, respecting the original definition for rarefaction. Rarefying is most often defined by the following steps (13).
>
> 1. Select a minimum library size, N_L,m_. This has also been called the rarefaction level (26), though we will not use the term here.
> 2. Discard libraries (microbiome samples) that have fewer reads than N_L,m_.
> 3. Subsample the remaining libraries without replacement such that they all have size N_L,m_.
>
> Often N_L,m_ is chosen to be equal to the size of the smallest library that is not considered defective, and the process of identifying defective samples comes with a risk of subjectivity and bias. In many cases researchers have also failed to repeat the random subsampling step (3) or record the pseudorandom number generation seed/process — both of which are essential for reproducibility.”

It was unfortunate that McMurdie and Holmes used the term “rarefying” in this quote and throughout their manuscript. The authors were correct to state that the distinction between “rarefying” and “rarefaction” is confusing and leads to their conflation. Adding to the confusion is that the papers cited in the first sentence of this quote (i.e., (13, 26, 27)) either do not use the words “rarefy” or “rarefying” or use them interchangeably with “rarefaction”. For example, Hughes and Hellman did not use “rarefy” and use “rarefaction” in the traditional sense with multiple subsamplings (13). Meanwhile, the QIIME-based literature appears to use “rarefy” and “rarefaction” interchangeably to mean only a single subsampling (26, 27). Subsequent researchers have continued to conflate the terms when citing WNWN (**Supplemental Text**). An exemplar of the confusion is the creation of a technique that uses “repeatedly rarefying” as an approach distinct from rarefaction when they were re-proposing traditional rarefaction (30).

As shown in the quoted text from WNWN, McMurdie and Holmes did emphasize the distinction between rarefying and rarefaction. However, because they seem to have coined a new definition for rarefying, they added to the confusion by using the generally used verb form of rarefaction. Further confusion comes from the author’s admonition in the final sentence of the above quoted text that some researchers have failed to repeat the subsampling step. Traditionally, repeating the subsampling step a large number of times and averaging the result is rarefaction (12, 13, 28). In other words, rarefying or subsampling is rarefaction, but with a single randomization. To minimize confusion, I will use “subsampling” in place of “rarefying” through the remainder of this study.

For clarity, I will use the following definition of rarefaction:

1. Select a minimum library size, N_L,m_. Researchers are encouraged to report the value of N_L,m_.
2. Discard samples that have fewer reads than N_L,m_.
3. Subsample the remaining libraries without replacement such that they all have size N_L,m_.
4. Compute the desired metric (e.g., richness, Shannon diversity, Bray-Curtis distances) using the subsampled data
5. Repeat steps 3 and 4 a large number of iterations (typically 100 or 1,000). Researchers are encouraged to report the number of iterations.
6. Compute summary statistics (e.g., the mean) using the values from step 4.

This definition aligns with how rarefaction was originally defined for comparing richness (i.e., the number of taxa in a community) across communities when communities are sequenced to different depths (5, 6). With this more general approach to the definition of rarefaction, rarefaction can be performed using any alpha or beta diversity metric. This strategy has been widely used by my research group and others and is available in the mothur software package using commands such as summary.single, rarefaction.single, phylo.diversity, and dist.shared. The procedure outlined above could also be used for hypothesis tests of differential abundance; however, consideration is needed to synthesize the results of these tests across a large number of replications.

## Description and critique of “Simulation A” from WNWN

McMurdie and Holmes analyzed the effect of subsampling and other approaches on clustering accuracy using what they called “Simulation A” in their Figure 2A and elsewhere in WNWN. In Simulation A, they investigated the ability to correctly assign simulated microbiome samples to one of two clusters representing two simulated treatment groups. Thankfully, the R code for Simulation A was provided by the authors in the simulation-cluster-accuracy/simulation-cluster-accuracy-server.Rmd R-flavored markdown (R markdown) file that was published as Protocol S1 in WNWN. I will outline the simulation strategy below and will reference line numbers from their R markdown file with “L” as a prefix.

**Figure 1.**
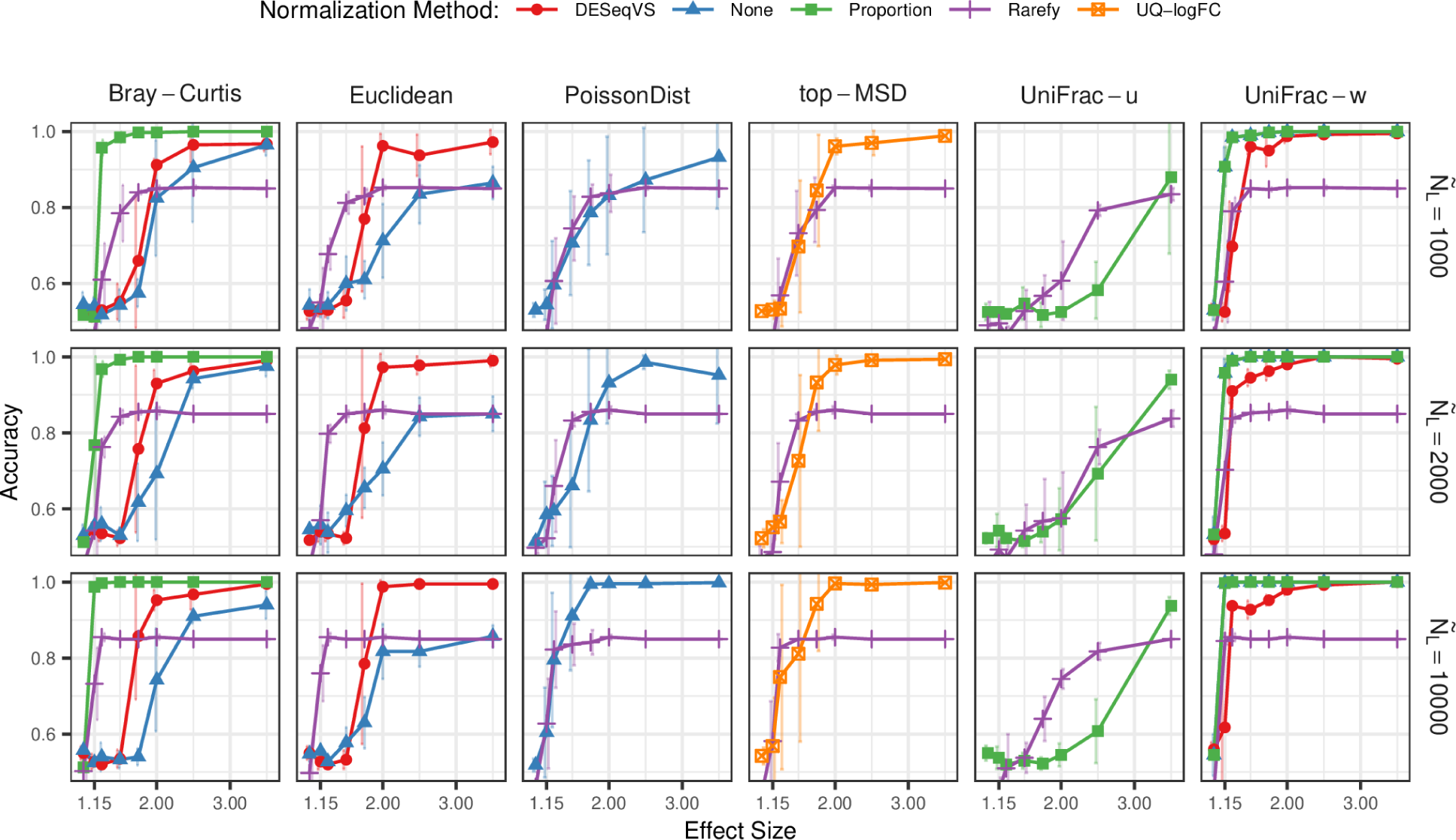
Re-running the R markdown provided in Protocol S1 of WNWN qualitatively reproduced Figure 4 from WNWN.

**Figure 2.**
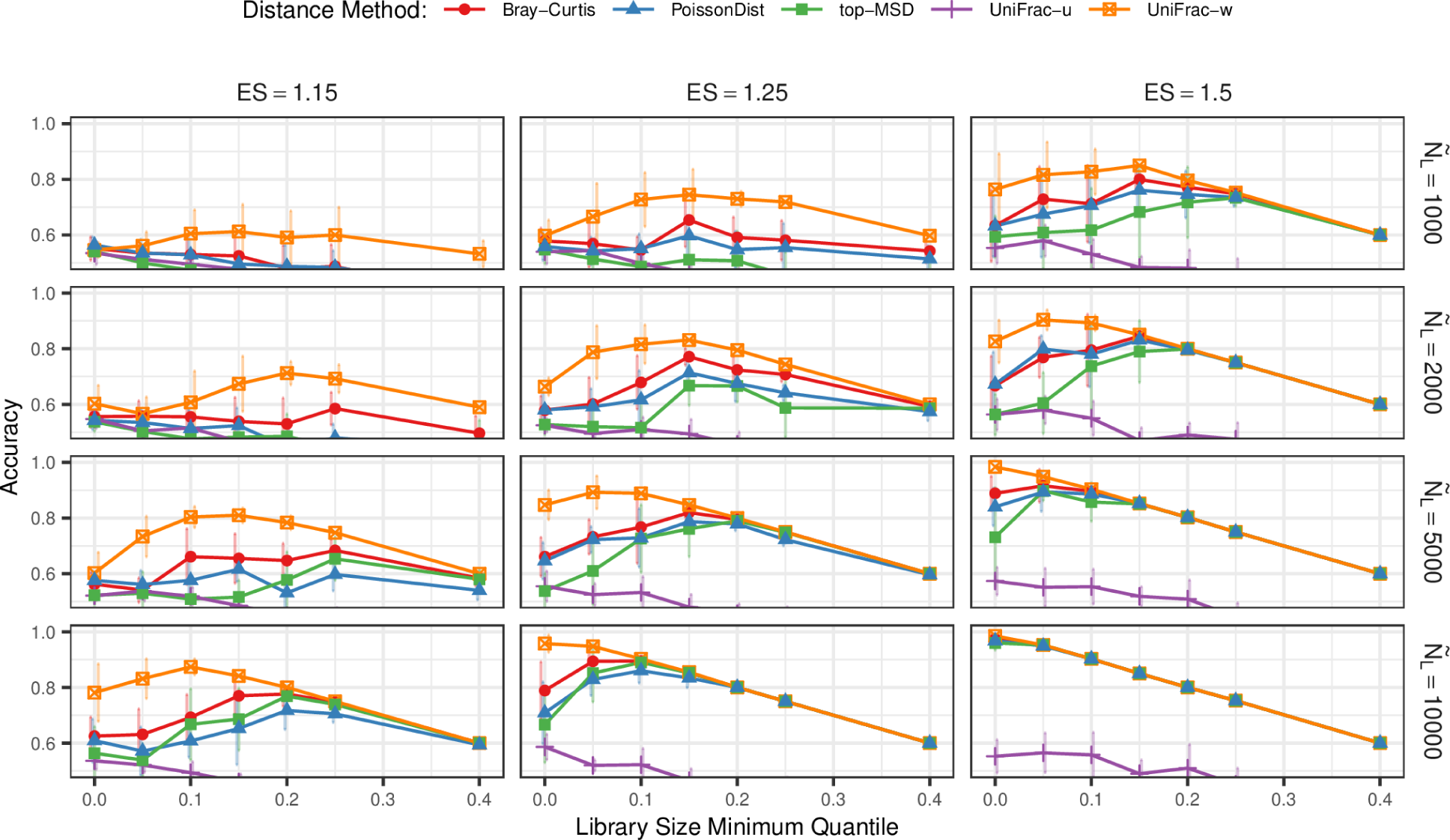
Re-running the R markdown provided in Protocol S1 of WNWN qualitatively reproduced Figure 5 from WNWN.

For the WNWN cluster analysis, OTU abundance distributions for the two treatment groups were generated using human fecal and ocean data originally take from the GlobalPatterns dataset (L129) (2). To generate the fecal and ocean distributions, the authors included any operational taxonomic unit (OTU) that appeared in more than one of the 4 fecal and 3 ocean samples (L60 and L137). The OTUs were sorted by how many of the 7 samples the OTUs were observed in followed by their total abundance across all 7 samples (L139). From this sorted list they obtained the identifiers of the first 2,000 OTUs (L66). Returning to the 7 samples they selected the data for the corresponding 2,000 OTUs and pooled the OTU abundances of the fecal and ocean samples separately to create two OTU abundance distributions (L144, L159-160, L197-198). Next, the fecal and ocean distributions were mixed in 8 different fractions to generate two community types that differed by varying effect sizes (i.e., 1.00, 1.15, 1.25, 1.50, 1.75, 2.00, 2.50, and 3.50; L170-195, L220); an effect size of 1.00 generated a null model with no difference between the treatment groups. To simulate the variation in sequencing depth across the 80 samples, they normalized the number of sequences from each of the 26 samples in the GlobalPatterns dataset so that the median number of sequences (Ñ_L_) across the samples had 1,000, 2,000, 5,000, or 10,000 sequences (L324-325). They then randomly sampled the 26 normalized sequencing depths, with replacement, to generate 80 sequencing depths. From each community type, they simulated 40 samples by sampling to the desired number of sequences (L73, L230-233 and L326-327). Each simulation condition was repeated 5 times (L85). Finally, they removed rare and low prevalence OTUs in two steps. First, they removed any OTUs whose total abundance was less than 3 across all 80 samples and that did not appear in at least 3 samples (L368-386). Second, they removed any OTUs that did not have more than 1 sequence in more than 5% of the 80 samples (i.e., 4 samples) and that did not have a total abundance across the 80 samples greater than one half of the number of samples in each community type (i.e., 20) (L523-538, L551). Simulation A consisted of 160 simulations (8 effect sizes x 4 median sequencing depths x 5 replicates = 160 simulations).

After generating the OTU counts for the 160 simulations, the authors applied several normalization methods, distance calculations, and clustering algorithms to the data. To normalize the OTU data the WNWN analysis either applied no normalization procedure (L559), calculated OTU relative abundances (L392-401, L562-563), subsampled the data (L409-416, L566-569), performed Variance Stabilization using the DESeq R package (L478-504, L580-583)(7), or performed Upper-Quartile Log-Fold Change normalization using the edgeR R package (L425-457, L576-577, L622-721) (19). In subsampling the data, the authors either included all of the samples or removed samples whose sequencing depth fell below the 5, 10, 15, 20, 25, and 40 percentile across all 80 samples (L96, L1065-1077).

Non-normalized data were used to calculate Bray-Curtis, Euclidean, Poisson, and Weighted UniFrac distances (L793-798, L801-806). Relative abundance data were used to calculate Bray-Curtis, Unweighted UniFrac, and Weighted UniFrac distances (L760-767). Subsampled data were used as input to calculate distances between samples using Bray-Curtis, Euclidean, Unweighted UniFrac, Weighted UniFrac distances, Poisson distances, and top-Mean Squared Difference (L769-775, L793-798, L801-806). DESeq Variance Stabilization normalized data were used to calculate Bray-Curtis, Euclidean, and Weighted Unifrac distances (L793-798). The Upper-Quartile Log-Fold Change normalized data were only used to calculate top-Mean Squared Difference distances (L801-806). WNWN calculated Bray-Curtis (31), Euclidean (31), Unweighted UniFrac (32), and Weighted UniFrac distances (33) using the Phyloseq R package (34). They calculated Poisson distances using the PoiClaClu R package (35). The top-mean squared difference was calculated using the edgeR R package (19). The resulting distance matrices were used to cluster the 80 samples into one of two clusters using partitioning around the medoid (PAM), K-means clustering, and hierarchical clustering (L865-879). Although data for all three methods were presented in Protocol S1, only the PAM data were presented in the main manuscript. The accuracy of the clustering assignments was quantified as the fraction of the 80 samples that were assigned to the correct cluster (L887-908). Since some of the subsampling conditions removed samples below a minimum sequencing depth threshold, those samples were counted as mis-clustered samples yielding minimum accuracies below 50%.

Although all simulations represent an artificial representation of reality and can be critiqued, I have identified eleven elements of the design of Simulation A that warranted further review.

1. Simulated conditions were only replicated 5 times each, potentially increasing the sensitivity of results to the choice of the random number generator seed
2. The average sizes of the libraries were small by modern standards, which may limit the generalizability of the results
3. DESeq-based Variance Stabilization generates negative values and was used with distance calculation methods that are sensitive to negative values, which likely led to nonsense distances and clusters
4. A single subsampling of each dataset was evaluated rather than using rarefaction, which likely resulted in noisier data
5. Results using PAM clustering were not directly compared to those of K-means and hierarchical clustering, although close inspection suggests that K-means may have been superior to PAM for some conditions
6. Subsampling removed the smallest 15% of the samples, which penalized accuracy values by 15 percentage points
7. The distribution of library sizes was not typical of those commonly seen in microbiome analyses, which may limit the generalizability of the results
8. A filtering step was applied to remove rare taxa from the simulated datasets, which may have skewed the shape of the community distributions
9. There was no accounting for differences in performance when library sizes are confounded with treatment group
10. Clustering accuracy was used rather than the more direct and frequently applied comparisons of beta diversity using permutation tests
11. There was no consideration of effects of normalization methods on alpha diversity metrics, which is the traditional application of rarefaction

Below, I replicated the original WNWN simulations and evaluated these points to reassess whether subsampling or rarefaction are “inadmissible”.

## Results

### Replication of WNWN simulations and results

Before assessing the impact of the points I critiqued above, I attempted to replicate the results shown in Figures 4 and 5 of WNWN using the authors’ code. I created a Conda environment that used the R version and package versions that were as close as possible to those used in WNWN. It was necessary to patch the WNWN’s R markdown file to render the document to be compatible with the overall workflow of this study. I was able to generate figures similar to those presented as WNWN’s Figures 4 and 5; my results are shown in this study as Figures 1 and 2, respectively. The differences in results are likely due to differences in software versions and operating systems. It is also worth noting that the published versions of the two figures differ from those included in Protocol S1 within the rendered HTML file (simulation-cluster-accuracy/simulation-cluster-accuracy-server.html) and that the figure numbers are one higher in the paper than those generated by the R markdown file (i.e., Protocol S1 labels the figures as Figures 3 and 4 corresponding to the published Figures 4 and 5). Regardless of the differences, my results were qualitatively similar to that provided in WNWN’s Protocol S1.

**Figure 3.**
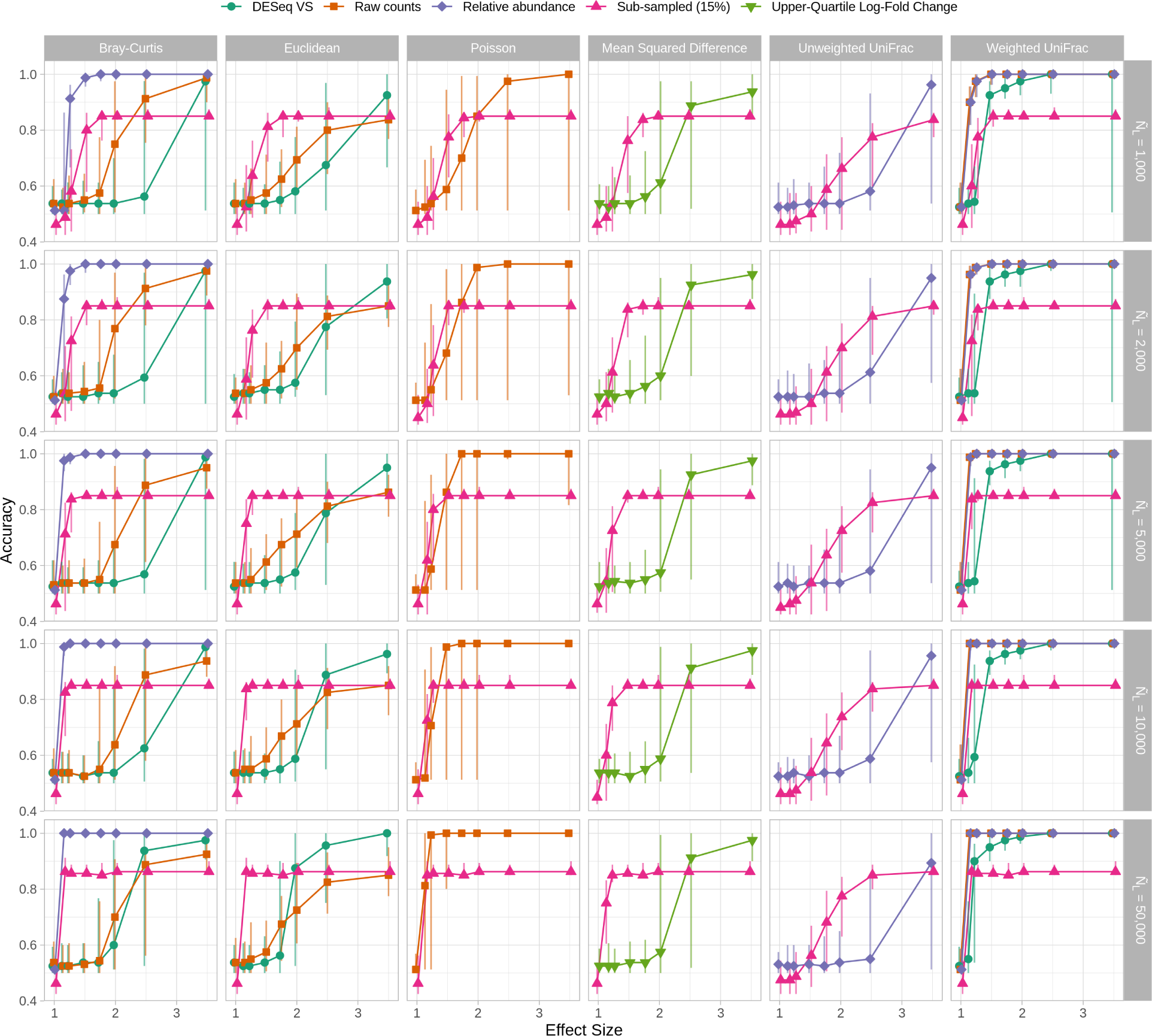
Successful reimplementation and expansion of analysis presented in Figure 4 from WNWN in a Snakemake pipeline. The reimplemented workflow largely borrowed from the original simulation-cluster-accuracy-server.Rmd R markdown file provided in WNWN’s Protocol S1. The most notable differences include the use of 100 rather than 5 randomizations and the addition of the median sequencing depth (Ñ_L_) of 50,000. The plotting symbols indicate the median of 100 randomizations and the error bars represent the observed 95% confidence interval. Simulations run at the same effect size are dodged to better reveal overlapping data. The sequencing depths were drawn from the GlobalPatterns dataset and sequences were clustered using PAM.

**Figure 4.**
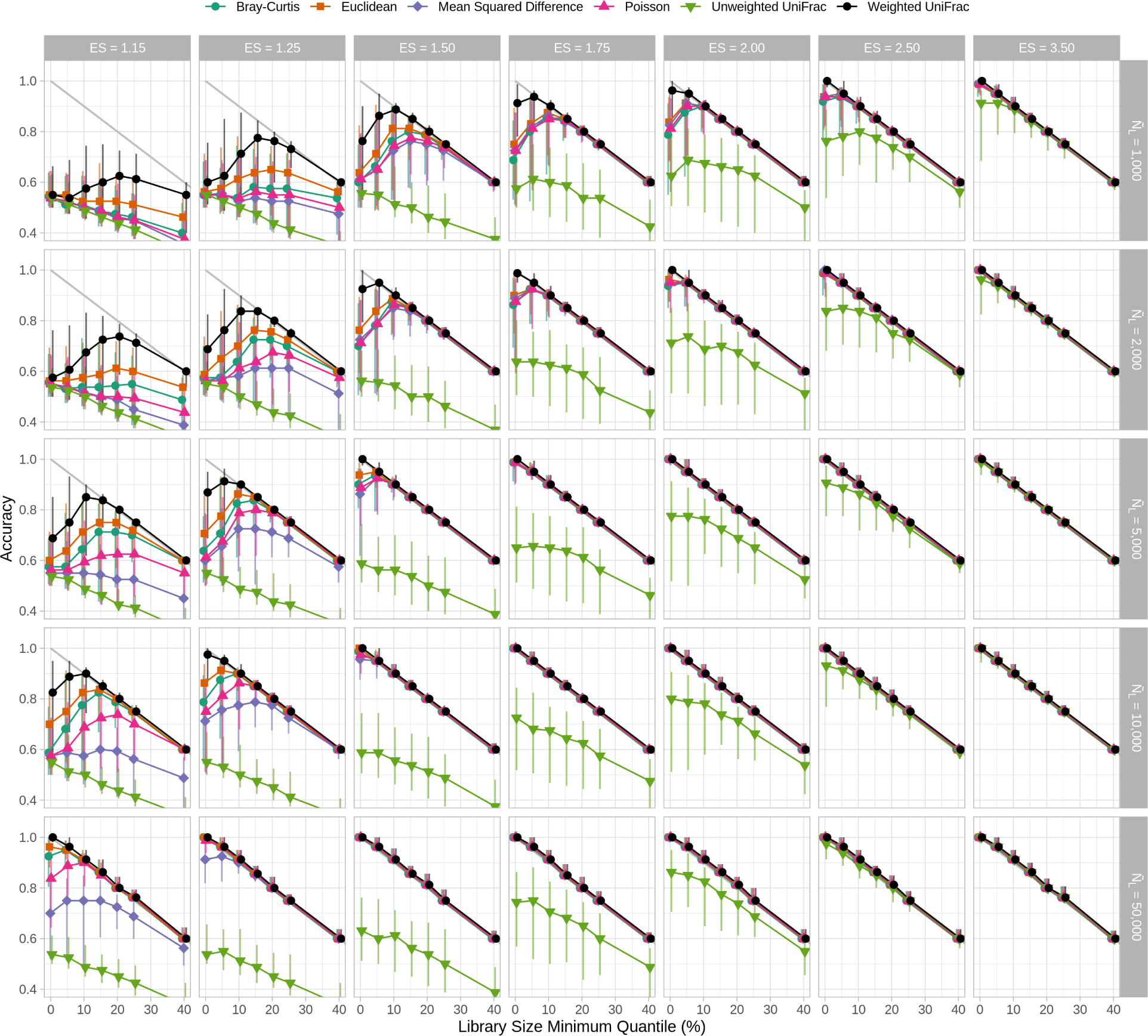
Successful reimplementation and expansion of analysis presented in Figure 5 from WNWN in a Snakemake pipeline. The reimplemented workflow largely borrowed from the original simulation-cluster-accuracy-server.Rmd R markdown file provided in WNWN’s Protocol S1. The most notable differences include the use of 100 rather than 5 randomizations and the addition of the median sequencing depth (Ñ_L_) of 50,000. The plotting symbols indicate the median of 100 randomizations and the error bars represent the observed 95% confidence interval. Simulations run at the same effect size are dodged to better reveal overlapping data. A light gray line is shown to indicate the best possible accuracy for each library size minimum quantile value. The sequencing depths were drawn from the GlobalPatterns dataset and sequences were clustered using PAM.

**Figure 5.**
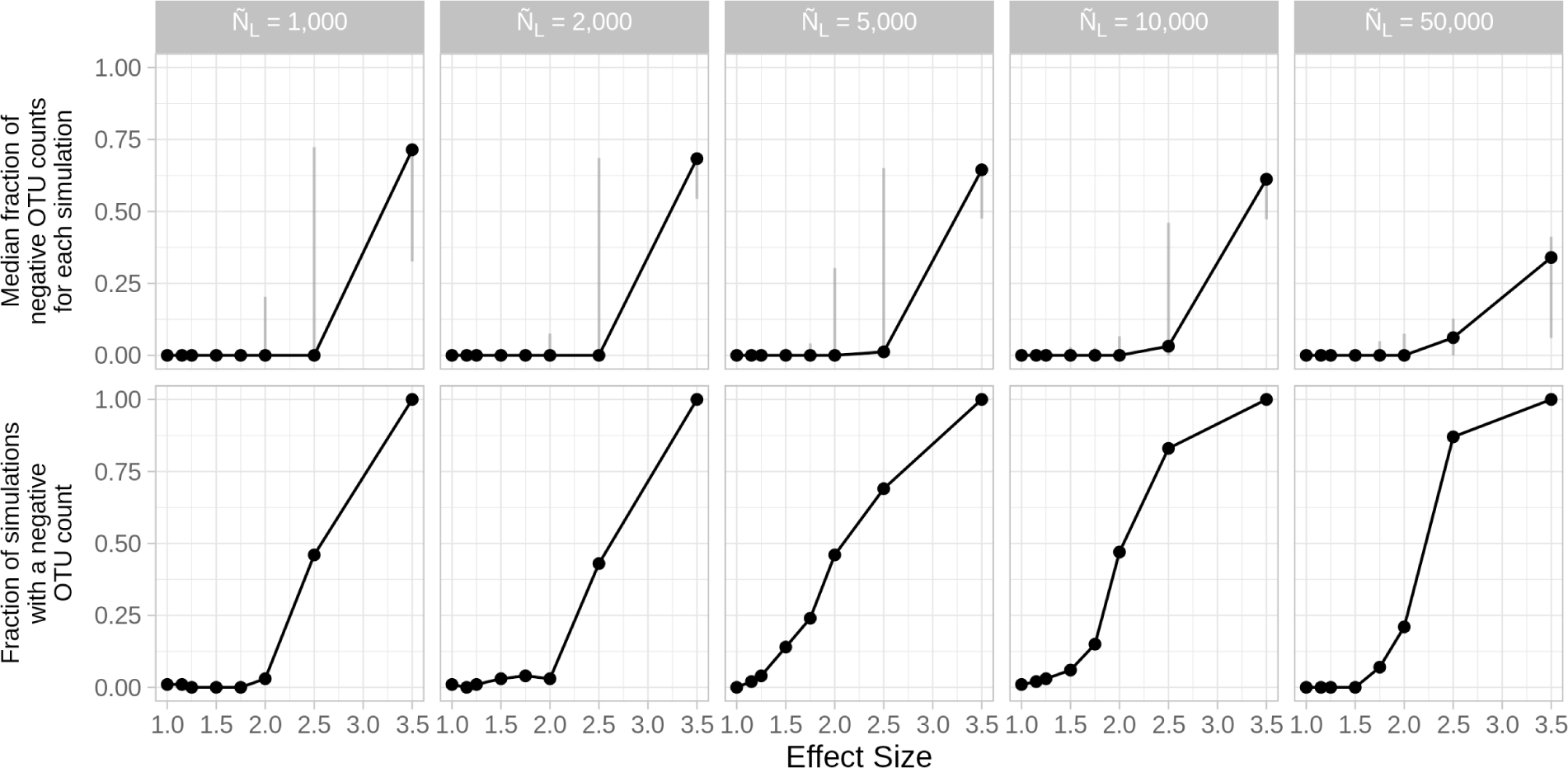
DESeq Variance Stabilization of OTU counts resulted in negative values that were used to calculate Bray-Curtis and Weighted UniFrac distances. The median number of negative OTU counts that had a negative OTU count following normalization increased with the effect size and decreased as Ñ_L_ increased (first row). The error bars indicate the observe 95% confidence interval. The fraction simulated datasets that had a negative OTU count following normalization also increased with effect size, but increased as Ñ_L_ increased (second row). For each effect size there were 100 replicate datasets.

### 1. Simulated conditions were only replicated 5 times each, potentially increasing the sensitivity of results to the choice of the random number generator seed

Each simulated condition was replicated 5 times in WNWN and the paper reported the mean and standard deviation of the replicate clustering accuracies. The relatively small number of replicates accounts for the jerkiness of the lines in WNWN’s Figures 4 and 5 (e.g. the Bray-Curtis distances calculated on the DESeqVS normalized data). A better approach would have been to use 100 replicates as this would reduce the dependency of the results on the random number generator’s seed. By increasing the number of replicates it was also possible to compare the probability of falsely and correctly clustering samples from the same and different treatment groups together (see points 10 and 11, below). Because the accuracies were not symmetric around the mean accuracy values the median and 95% confidence intervals or interquartile range should have been reported. To test the effect of increasing the number of replicates, I pulled apart the code in simulation-cluster-accuracy/simulation-cluster-accuracy-server.Rmd into individual R and bash scripts that were executed using a Snakemake workflow with the same Conda environment I used above. This was necessary since the number of simulated conditions increased 20-fold with the additional replicates. Such intense data processing was not practical within a single R markdown document. Again, the observed results were qualitatively similar to those generated using the single R markdown file (Figures 3 and 4). The increased number of replications resulted in smoother lines and allowed me to present empirical 95% confidence intervals. For all analyses in the remainder of this paper, I used 100 randomized replicates per condition.

### 2. The average sizes of the libraries were small by modern standards, which may limit the generalizability of the results

In the 10 years since WNWN was published, sequencing technology has advanced and sequence collections have grown considerably. For more modern datasets, it would be reasonable to expect a median number of sequences larger than 10,000 (see Table 1 of (4)). Therefore, I included an additional median depth of sequencing of 50,000 sequences with WNWN’s four median sequencing depths (i.e., 1,000, 2,000, 5,000, 10,000). Additional sequencing coverage would be expected to result in more robust distance values since there would be more information represented in the data. Indeed, the added sequencing depth showed higher accuracy values at lower effect sizes for the combinations of normalization methods and distance calculations (Figure 3). Increased sequencing coverage also resulted in improved clustering accuracy for lower effect sizes when the library size minimum quantile was decreased (Figure 4). I will revisit the choice of the library size minimum quantile below.

### 3. DESeq-based Variance Stabilization generates negative values and was used with distance calculation methods that are sensitive to negative values, which likely led to nonsense distances and clusters

Close comparison of WNWN’s Figure 4 and my version (Figure 3) revealed several difference between the two plots. First, in the WNWN analysis, the accuracies for the Weighted UniFrac distances at the largest effect size (i.e., 3.5) were 1.00 for median sequencing depths of 1,000, 2,000, and 10,000. In my version of the analysis, the values for the same sequencing depths were 1.00, 1.00, and 1.00, respectively. The 95% confidence interval for these accuracies spanned between 0.51 and 1.00. Second, clustering accuracy for Bray-Curtis and Weighted UniFrac distances calculated using DESeq Variance Stabilization normalized OTU counts were different between the original and my simulation at smaller effect sizes and had wide confidence intervals. Inspection of the DESeq Variance Stabilization normalized OTU counts revealed that the method resulted in negative values. It has been suggested that WNWW turned negative DESeq normalized counts to zero (25); however, I was unable to find code in simulation-cluster-accuracy-server.Rmd that made this transformation. In fact, rendering the R markdown files in WNWN’s Protocol S1 generated warning messages when passing the DESeq normalized counts to the Bray-Curtis calculator, which said, “results may be meaningless because data have negative entries in method ‘bray’”. Although the Weighted UniFrac calculator function did not generate a similar warning message, negative count values would also result in similarly meaningless distances. Both are due to the fact that the distance calculators sum the counts of each OTU in both samples being compared. In contrast, a Euclidean distance does not use a similar sum, but sums the square of the difference between the OTU abundance in each sample. Even if negative values were converted to zeroes, this would effectively the same as removing rare taxa, which could have a significant impact on the shape of the communities (4). To assess the prevalence of negative counts in the simulated data, I quantified the fraction of negative values in the OTU matrix from each simulation and counted the number of simulations where the normalized OTU table had at least one negative value (Figure 5). In general the fraction of negative OTU counts increased with effect size, but decreased with sequencing effort. The fraction of simulations with at least one negative value increased with effect size and sequencing effort. The high frequency of negative OTU counts resulted in highly variable Bray-Curtis and Weighted UniFrac values. It is likely that because the WNWN analysis only used 5 replicates that the large variation in accuracies at high effect sizes was missed initially. For the rest of this reanalysis study, I will only report results using the DESeq-based Variance Stabilization normalization with the Euclidean distance.

### 4. A single subsampling of each dataset was evaluated rather than using rarefaction, which likely resulted in noisier data

A more robust analysis would have used rarefaction since it would have averaged across a large number of random subsamplings (e.g., 100 or 1,000). By using a large number of subsamplings the likelihood of incorporating all of the OTUs would have increased. Rather than being guilty of “omission of available valid data” as claimed in WNWN, with a sufficient number of subsamplings, traditional rarefaction uses all of the available data. To fairly compare subsampling, as employed in WNWN, and rarefaction, I removed the 15% of samples with the lowest number of sequences and compared the clustering accuracies from a single subsampling to rarefaction with 100 randomizations. This analysis revealed two benefits of rarefaction. First, the median distances generated by rarefaction was always at least as large as those from a single subsample (Figure 6). The difference was most pronounced for smaller average library sizes and at smaller effect sizes. The Unweighted UniFrac distances were most impacted by the use of rarefaction over subsampling. Second, the interquartile ranges in clustering accuracy by rarefaction were generally smaller than those by subsampling and showed similar trends to the difference in the median distances (Figure 6). Because rarefaction incorporates more of the data and generally performed better than subsampling, the remainder of this analysis will report results using rarefaction rather than by subsampling, except when noted.

**Figure 6.**
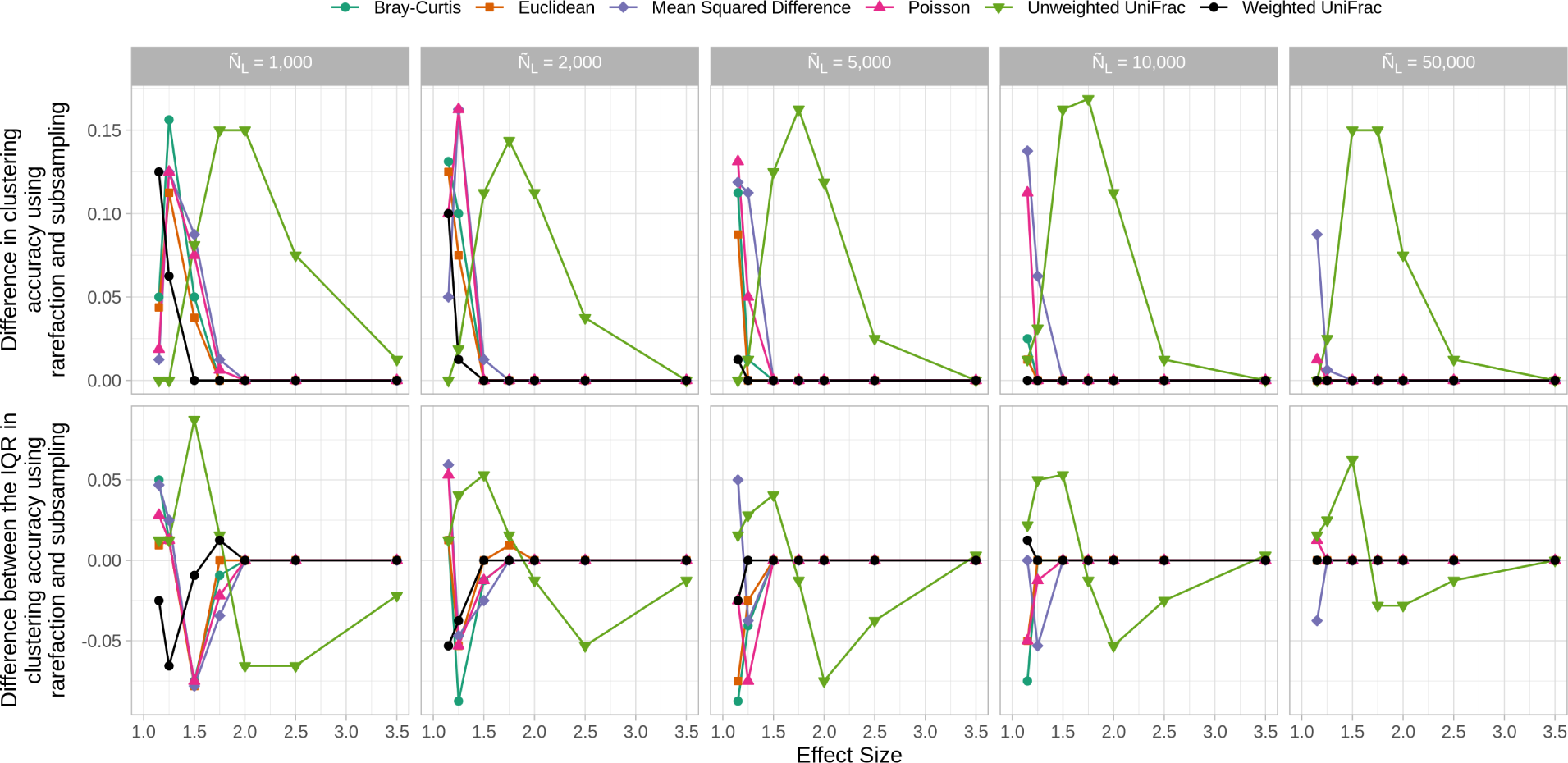
Rarefaction resulted in larger and less variable clustering accuracies. With the exception of Unweighted UniFrac distances, the improved performance by rarefaction was observed at smaller effect sizes. In the first row of panels larger values mean that the accuracies by rarefaction were better than those of subsampling. In the second row of samples, larger values mean that interquartile range (IQR) for rarefaction was larger than that of subsampling.

### 5. Results using PAM clustering were not directly compared to those of K-means and hierarchical clustering, although close inspection suggests that K-means may have been superior to PAM for some conditions

The clustering accuracy measurements reported in the body of WNWN were determined using PAM-based clusters while Protocol S1 also includes K-means and hierarchical clustering. Although the data were not displayed in a manner that lent itself to direct comparison in Protocol S1, close inspection of the rendered figures suggested that PAM may not have been the optimal choice in all situations. Actually, K-means clustering appeared to perform better in many simulations. Because the accuracies were the smallest at lower effect sizes, I focused my comparison at the effect size of 1.15. For each set of 100 replicated simulated datasets, I compared the clustering accuracy across clustering methods to see how often each clustering method resulted in the highest accuracy (Figure 7). Indeed, K-means clustering performed better than the other methods. Among all combinations of normalization methods, distance calculations, and read depths, K-means clustering was at least as good as the other methods in 74.39% of the randomizations (Figure 7); PAM clustering resulted in clustering accuracies as good or better than the other methods in 49.92% of the randomizations; and hierarchical clustering (HClust) was at least as good as the other methods for 44.32% of the randomizations. Finally, I specifically compared the clustering accuracies using rarefaction for each of the distance calculations methods using PAM and K-means clustering. Among the 30 combinations of distance calculations and read depths, K-means performed better than PAM in 29 cases with PAM doing better in the 1 other case (i.e., calculating distances with Euclidean using 10,000 sequences). When using subsampled data, K-means clustering performed better than PAM in each case. Because K-means clustering did so much better than PAM clustering in the simulated conditions, I used K-means clustering for the remainder of this study.

**Figure 7.**
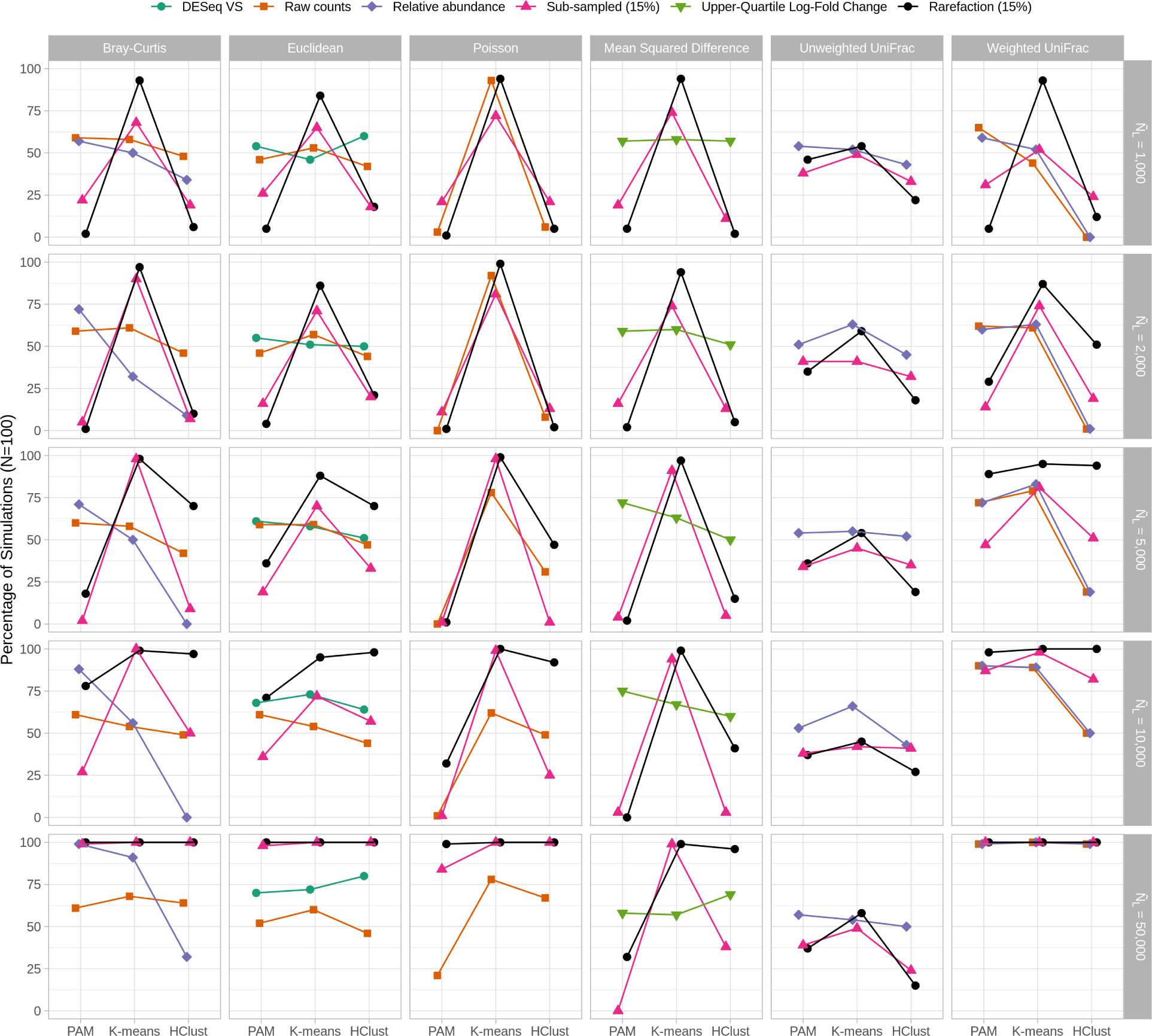
K-means clustering was consistently as good or better than PAM or hierarchical clustering when comparing rarefaction to other normalization methods. Each point represents the percentage of 100 simulations where that clustering method performed as well or better than the other methods for that normalization procedure.

### 6. Subsampling removed the smallest 15% of the samples, which penalized accuracy values by 15 percentage points

In WNWN, the authors quantified the tradeoff between median sequencing depth, the number of samples removed below the threshold, and clustering accuracy (WNWN’s Figure 5, my Figure 4). Although the optimal threshold varied by distance metric, normalization method, and sequencing depth, they removed samples whose number of sequences was less than the 15th percentile (L404-419). They acknowledged that this screening step, which was only used with subsampling, would decrease clustering accuracy putting it at a relative disadvantage to the other methods (page 5, column 1, last paragraph). Therefore, it was not surprising that the peak clustering accuracy for their subsampled data was at 85%. Because the true best threshold would not known *a priori* in an actual microbiome study, it would be impossible for researchers to conduct a sensitivity analysis comparing the tradeoffs between sequencing depth, sample number, and clustering accuracy to select a sequencing depth for their analysis without the risk of p-hacking. The differences in clustering accuracy between subsampling and rarefaction with PAM and K-means clustering indicated that it was necessary to reassess the tradeoff between the library size minimum quantile and clustering accuracy. When using rarefaction, K-means clustering, and only considering conditions with 2,000 or more sequences, there was not a condition where setting a higher threshold resulted in a better accuracy than using all of the samples (Figure 8). These results showed that for modern sequencing depths, using the full datasets with rarefaction and K-means clustering resulted in accuracies that were better than those observed when removing the smallest 15% of the samples from each simulated dataset. When the WNWN Figure 4 was recast with these approaches, rarefaction performed at least as well as any of the other transformations with each distance calculation (Figure 9).

**Figure 8.**
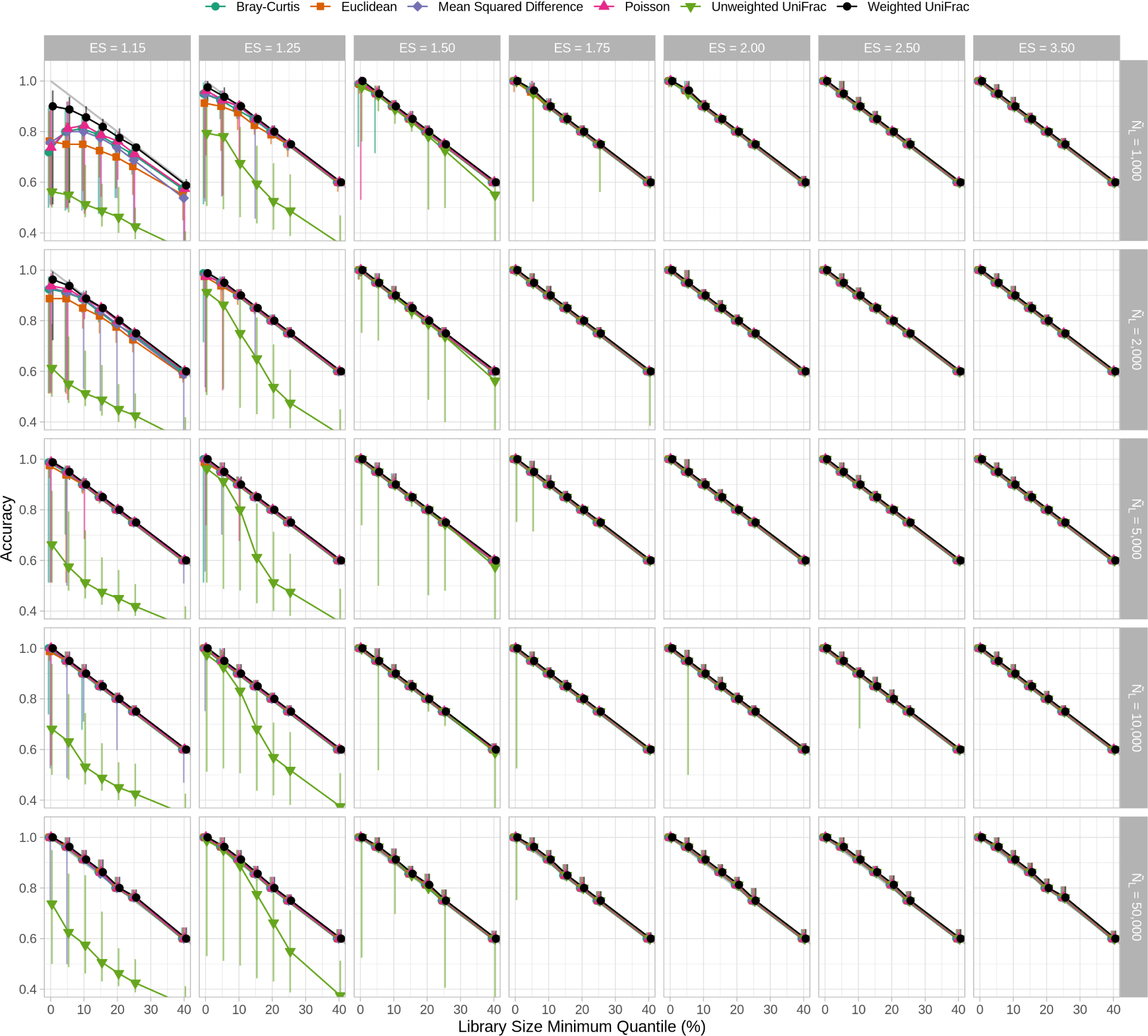
When the median sequencing depth was 2,000 sequences or more, rarefaction of the entire dataset performed better than removing the smallest 15% of samples when using K-means clustering. This figure is analogous to Figure 4 except that K-means clustering was used instead of PAM.

**Figure 9.**
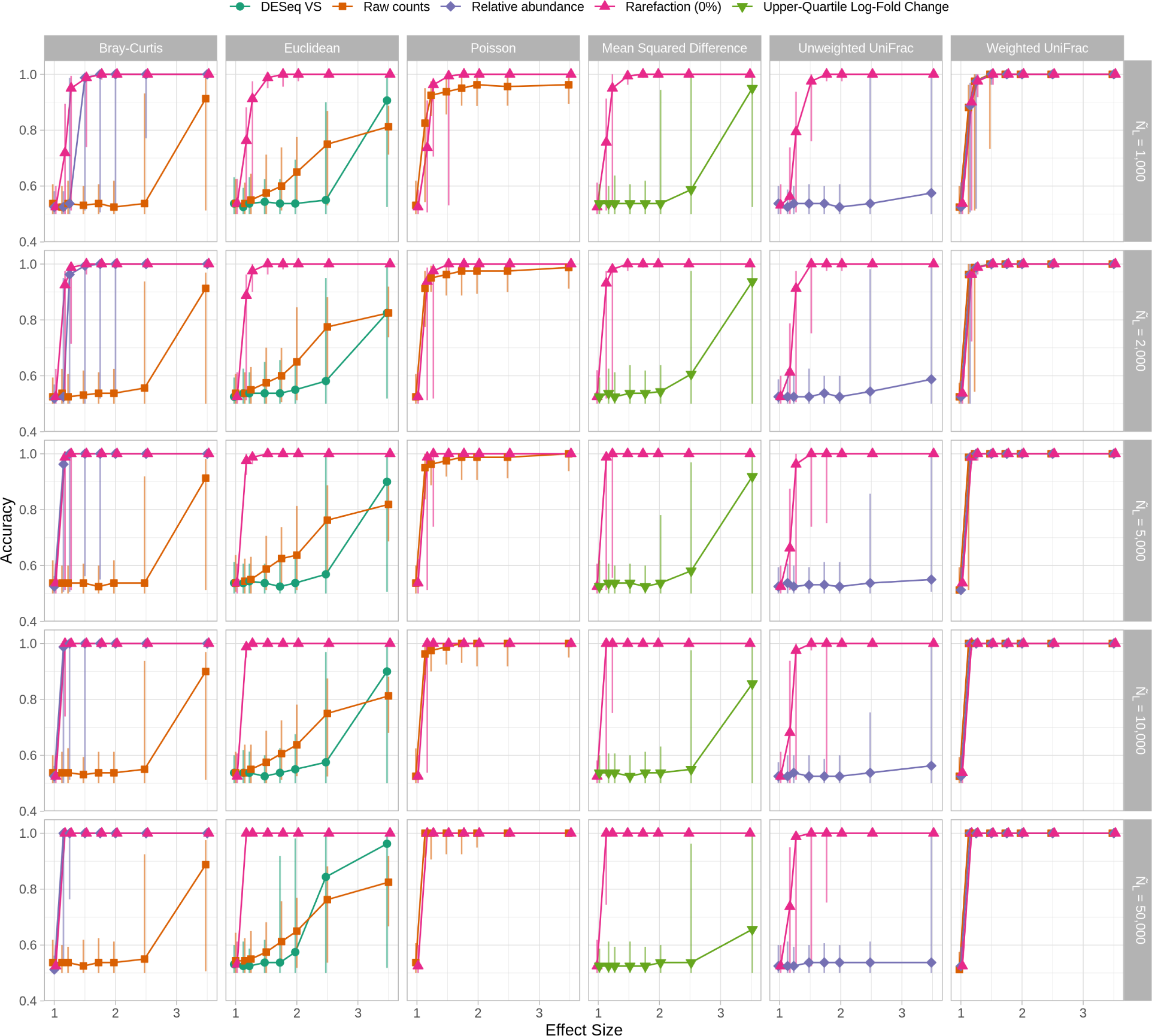
K-means clustering of distances calculated with rarefaction were as good or better than any other normalization method. This figure is analogous to Figure 3 except that K-means clustering was used instead of PAM, rarefaction on the full dataset was used instead of subsampling to the size of the sample at the 15th percentile, and DESeq Variance Stabilization normalized OTU counts were only used with Euclidean distances.

### 7. The distribution of library sizes was not typical of those commonly seen in microbiome analyses, which may limit the generalizability of the results

As described above, the sequencing depths used in the 26 GlobalPatterns datasets were used as the distribution to create sequencing depths for the 80 samples that were generated in each simulation. The GlobalPatterns datasets had a mean of 1,085,256.8 sequences and a median of 1,106,849 sequences per dataset (Figure 10). The datasets ranged in sequencing depth between 58,688 and 2,357,181 sequences for a 40.16-fold difference. Rather than representing a typically observed distribution of sequencing depths that would be skewed right (see Figure S1 from (4)), the sequencing depth distribution was normally distributed (Shapiro-Wilk test of normality, P=0.57). From these simulations it is unclear how sensitive the various normalizations and distance calculations were to a more realistic skewed distribution. A second limitation of this sequencing depth distribution is that it only contained 26 unique sequencing depths such that each sequencing depth would have been re-used an average of 3.08 times in each simulation. Yet, it is unlikely for a real sequence collection to have duplicate sequencing depths. To reassess the WNWN results in the context of a more typical distribution of sequencing depths, I created a new set of simulations to test the effect of the shape of the distribution on the results. I created a simple sequencing depth distribution where there were 80 depths logarithmically distributed between the minimum and maximum sequencing depths of the GlobalPatterns dataset (Figure 10). The median of this distribution was 372,040 and the mean was 629,824.8. When I regenerated the WNWN’s Figures 4 and 5 using the log-scaled sequencing effort distribution, the differences in normalization methods were more apparent (Figure 11 and 12). For each of the distance calculators, rarefaction to the size of the smallest dataset yielded accuracies that were at least as good as the other methods across effect sizes and median sequencing depths. The difference was most pronounced at smaller effect sizes and sequencing depths. When comparing the performance of rarefaction across distance calculators for different effect sizes, sequencing depths and size of smallest sample, the accuracies I observed using the log-scaled sample sizes was at least as good as those obtained using the GlobalPatterns-based distribution (Figure 9).

**Figure 10.**
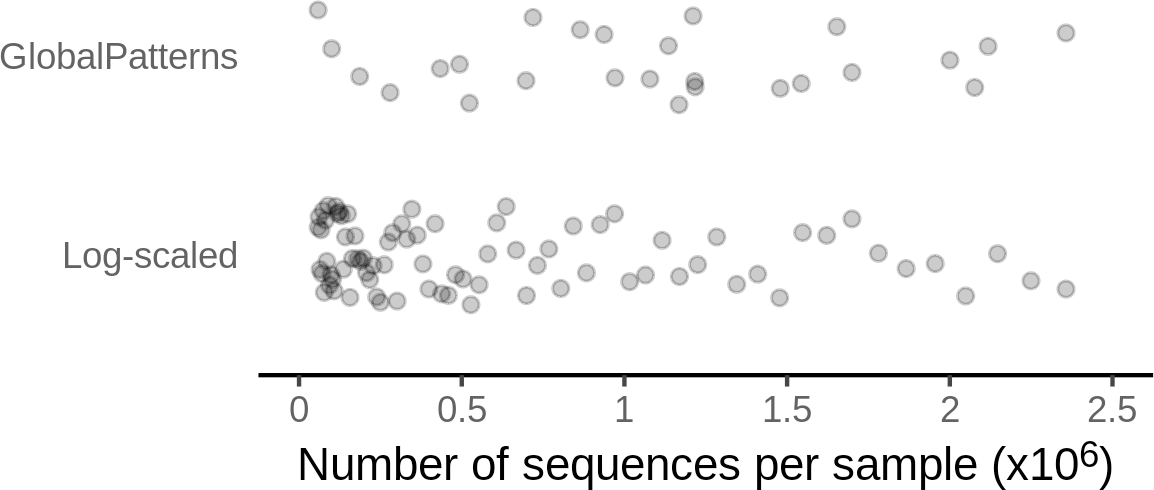
Comparison of the normally distributed sequencing depths from the GlobalPatterns dataset and a log-scaled distribution of sequencing depths. The log-scaled distribution was generated so that each sample in a simulation could have a unique number of sequences and to simulate the skew right distribution commonly seen in microbiome studies.

**Figure 11.**
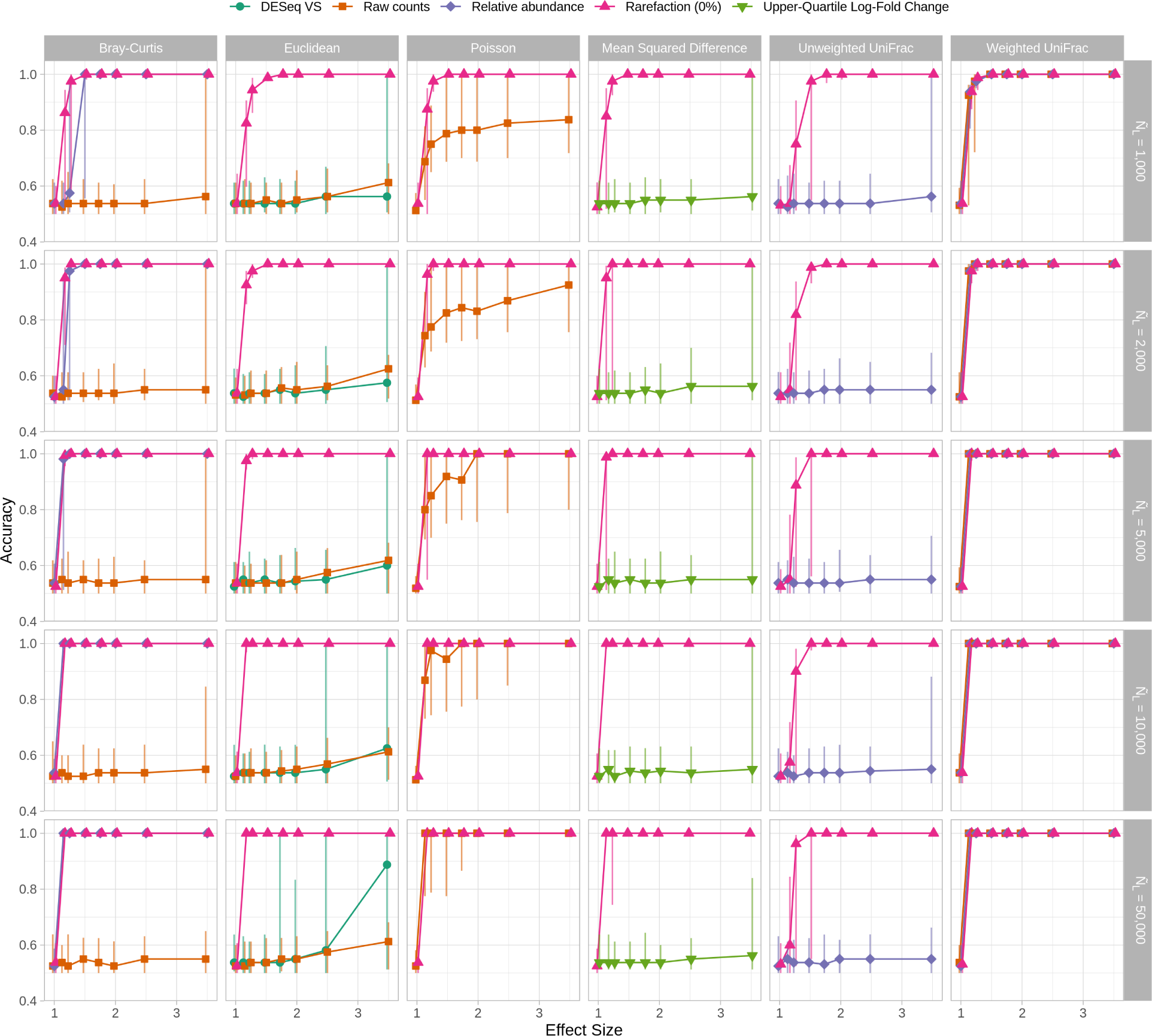
Clustering accuracies that used rarefaction were as good or better than the other normalization procedures when a log-scaled distribution of sequencing depths. This figure is analogous to Figure 9 except that the sequencing depths for each of the 80 samples in each simulation were drawn without replacement from a log-scaled distribution rather than from the GlobalPatterns sequencing depths.

**Figure 12.**
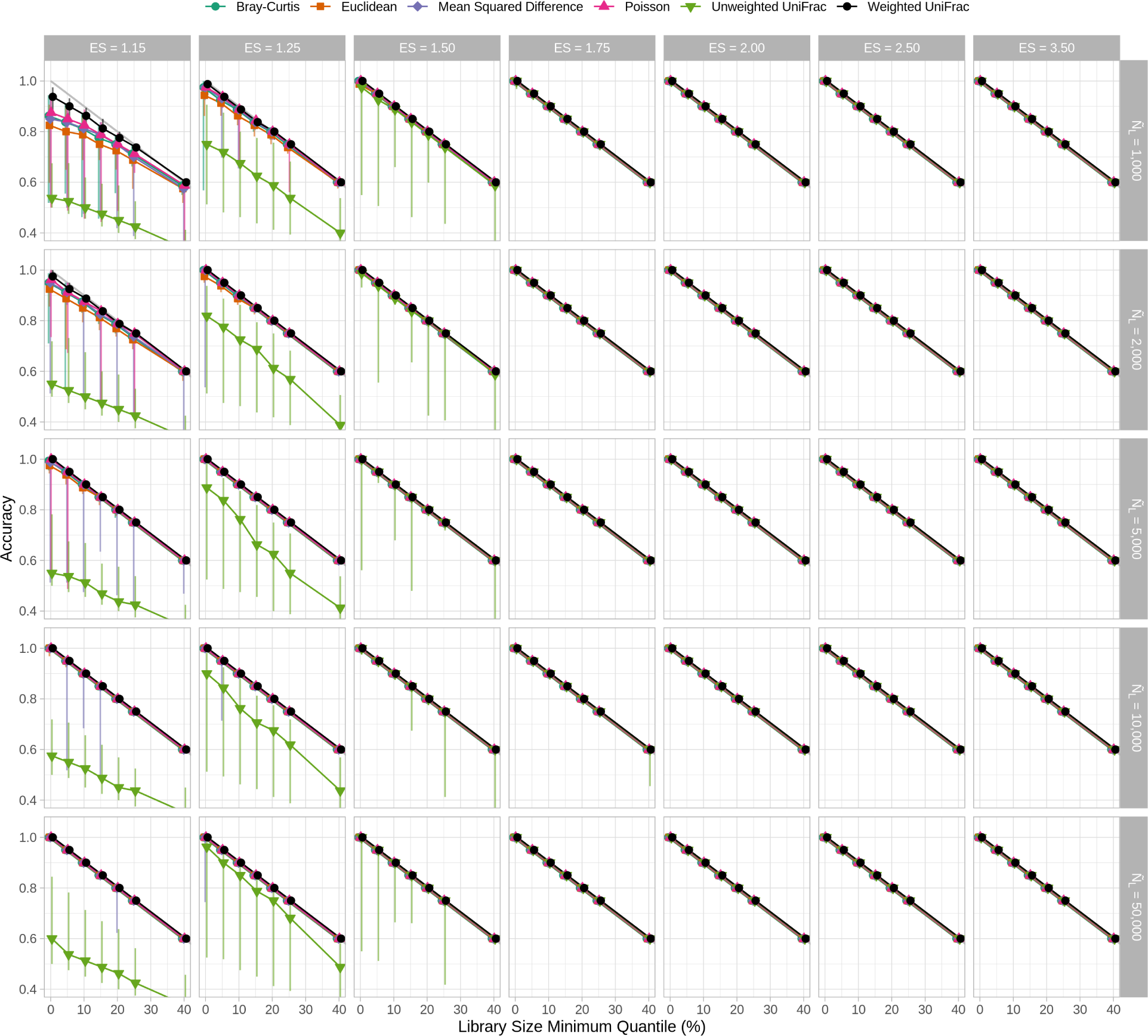
Rarefaction with all samples yielded clustering accuracies that were as good or better than removing the smallest 15% of samples across distance calculation methods. This figure is analogous to Figure 8 except that the sequencing depths for each of the 80 samples in each simulation were drawn without replacement from a log-scaled distribution rather than from the GlobalPatterns sequencing depths.

### 8. A filtering step was applied to remove rare taxa from the simulated datasets, which may have skewed the shape of the community distributions

McMurdie and Holmes were emphatic that “**rarefying biological count data is statistically inadmissible** because it requires the omission of available valid data” (emphasis in original). Thus it is strange that they argue against removing data when rarefying/subsampling, but accept removing rare and low-prevalence OTUs prior to normalizing their counts. This practice has become common in microbiome studies and is the standard approach in tools such as Dada2, Unoise, and Deblur (36–39). However, my previous work has shown that rare sequences from a poorly sequenced sample often appear in more deeply sequenced samples suggesting that they are not necessarily artifacts (4). Furthermore, removing rare sequences alters the structure of communities and has undesirable effects on downstream analyses. Although my previous work does an extensive analysis of the effects of removing rare sequences, I wanted to explore the effect of filtering in the context of the WNWN simulation framework. For each of the filtered and non-filtered OTU tables I calculated the the difference in accuracy between replicates for each normalization method and distance calculator across effect sizes for a Ñ_L_ of 10,000 (Figure 13). The median difference in accuracies (i.e. clustering accuracies with filtered data minus those without filtering) did not deviate meaningfully from zero. However, the 95% confidence intervals were most pronounced at large effect sizes when using raw counts, DESeq Variance Stabilization and Upper-Quartile Log-Fold Change and at smaller effect sizes when using rarefaction and relative abundances. Given the large variation caused by removing rare taxa and my previous work, OTU filtering should not be performed in microbiome analyses. Considering the minimal effect that removing rare OTUs had on the median difference in clustering accuracy in the current simulation framework, I have used the filtered datasets throughout the current study.

**Figure 13.**
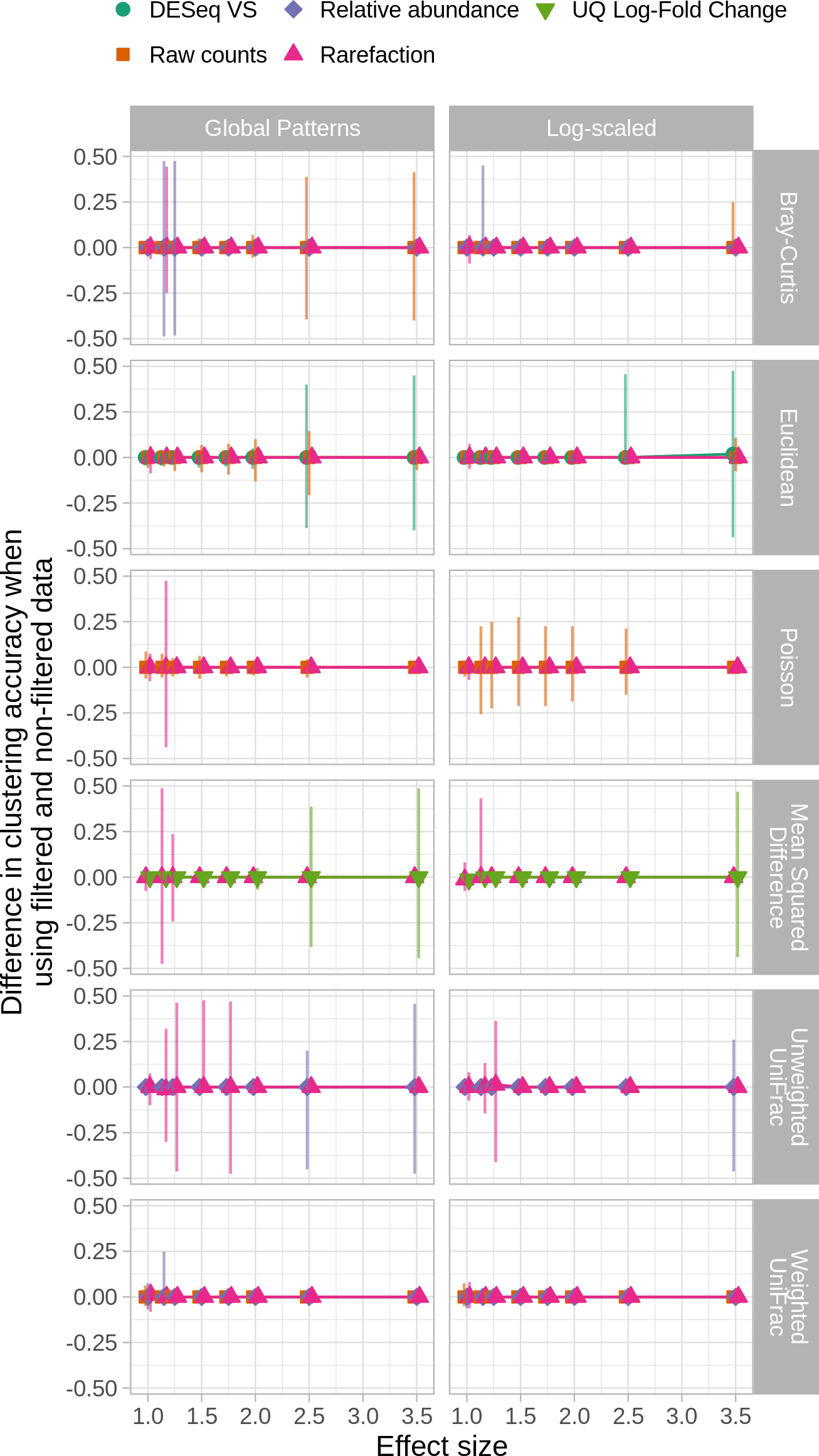
Normalization and distance calculation methods vary in their sensitivity to removal of rare OTUs. Larger values indicate that the clustering accuracy from filtered datasets were larger than those from non-filtered datasets. The median of 100 randomizations did not meaningfully vary from 0.0, but the observed 95% confidence interval varied considerably. Data are shown for a median sequencing depth (Ñ_L_) of 10,000 sequences when individual sequencing depths were sampled with replacement from the GlobalPatterns dataset or without replacement from the log-scaled distribution.

### 9. There was no accounting for differences in performance when library sizes are confounded with treatment group

I and others have observed that not using rarefaction can lead to falsely detecting differences between communities when sequencing effort is confounded with the treatment group (25, 40). Previous analyses showed that in these situations rarefaction did the best job of controlling the rates of false detection (i.e., Type I errors) and maintaining the statistical power to detect differences (i.e., 1-rate of Type II errors) of differences between groups of samples. Such situations have been observed when comparing communities at different body parts where one site is more likely to generate contaminating sequence reads from the host (e.g., 41). To determine whether this result was replicated with the WNWN simulation framework, I created a sequencing depth distribution where sequencing depth was fully confounded with the treatment group using both the GlobalPatterns and Log-scaled sequence distributions. To confound the sequencing depth, sequencing depths from one treatment group were drawn from below the median number of sequences of the sequencing distribution and those for the second treatment group were from above the median. To assess the risk of falsely detecting clusters, I compared the clustering accuracies using an effect size of 1.00 using both the confounded and randomized sequencing distributions (see rows 1 and 2 of Figure 14). The samples should have only been assigned to one cluster; however, each of the clustering methods forced the samples into two clusters. So, when there are two groups of 40 samples that do not differ, the best a method could do would be to correctly assign 41 of the 80 samples for an accuracy of 0.51. The false detection risk did not vary by method when the sequencing depth was randomized across treatment groups. Yet, when the sequencing depth was confounded with treatment group, rarefaction was the most consistent normalization method for controlling the generation of spurious clusters. At larger effect sizes the ability to correctly identify two clusters increased when the sequencing depth was confounded with treatment group (Figure 14). At the effect size of 1.15, the rarefied data generated the highest accuracy clusters regardless of whether the sequencing depths were confounded. Although the level of confounding in this simulation was extreme, it highlights the ability of rarefaction to control the false detection rate and the ability to correctly detect clusters relative to the other normalization methods.

**Figure 14.**
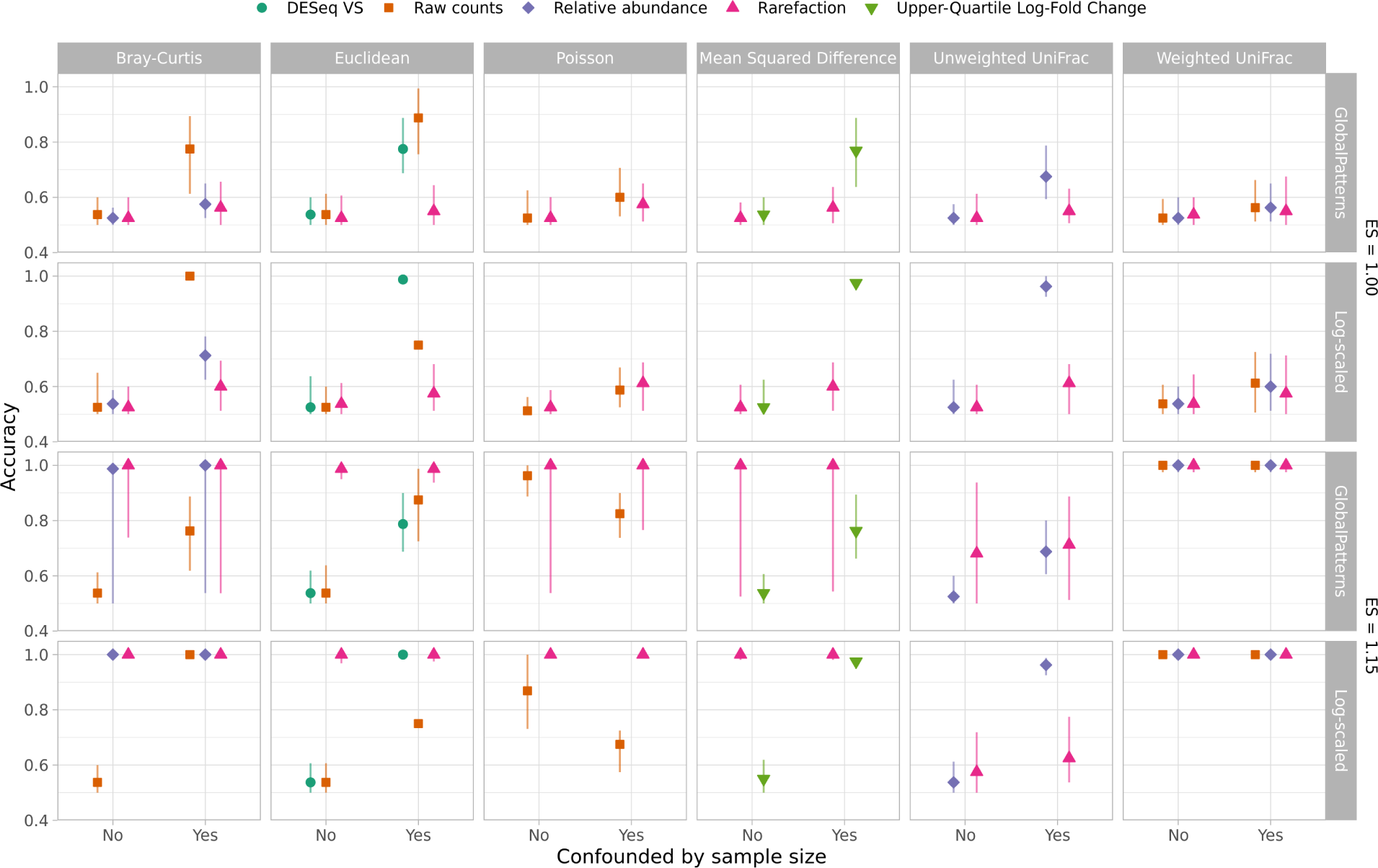
Rarefaction was consistently as good or better than all other normalization methods at assigning samples to the correct treatment group regardless of whether sequencing depth was confounded by treatment group. Because the clustering algorithms forced samples into one of two groups, the expected accuracy with an effect size of 1.00 was 0.51. With an effect size of 1.15, the expected accuracy was 1.00. Each point represents the median of 100 replicates and the error bars represent the observed 95% confidence interval. Data are shown for a median sequencing depth (Ñ_L_) of 10,000 sequences when individual sequencing depths were sampled with replacement from the GlobalPatterns dataset or without replacement from the log-scaled distribution.

### 10. Clustering accuracy was used rather than the more direct and frequently applied comparisons of beta diversity using permutation tests

Since WNWN was published, there has been controversy over the use of clustering methods to group samples (i.e., enterotypes). Concerns have been raised including whether such clustering should be done on ecological distances or sequence counts and the biological interpretation of such clusters (27, 42, 43). As described in the previous point, one notable challenge with using clustering accuracy as the dependent variable is that the clustering methods force the samples into one of two clusters. For the case where the effect size was 1.00, it was impossible for all 80 samples to be assigned to a single cluster. A more commonly used approach for analyzing distance matrices is to use a non-parametric analysis of variance test (i.e., AMOVA, PERMANOVA, NP-ANOVA)(44). I subjected each of the distance matrices to such a test using adonis2, a function from the vegan R package that implements this test to assess the effects of each normalization and distance calculation method on the Type I errors and statistical power (45). When sequencing depths were randomly distributed across the two treatment groups, the Type I error did not meaningfully deviate from the expected 5% (Figure 15). However, when sequencing depths were confounded with treatment group, rarefaction was the only normalization approach to control the Type I error. Similar to the clustering accuracy results, when distances were calculated using rarefaction, the tests consistently had the best statistical power (Figure 15). When considering both Type I error and power, rarefaction performed the best among the different normalizations.

**Figure 15.**
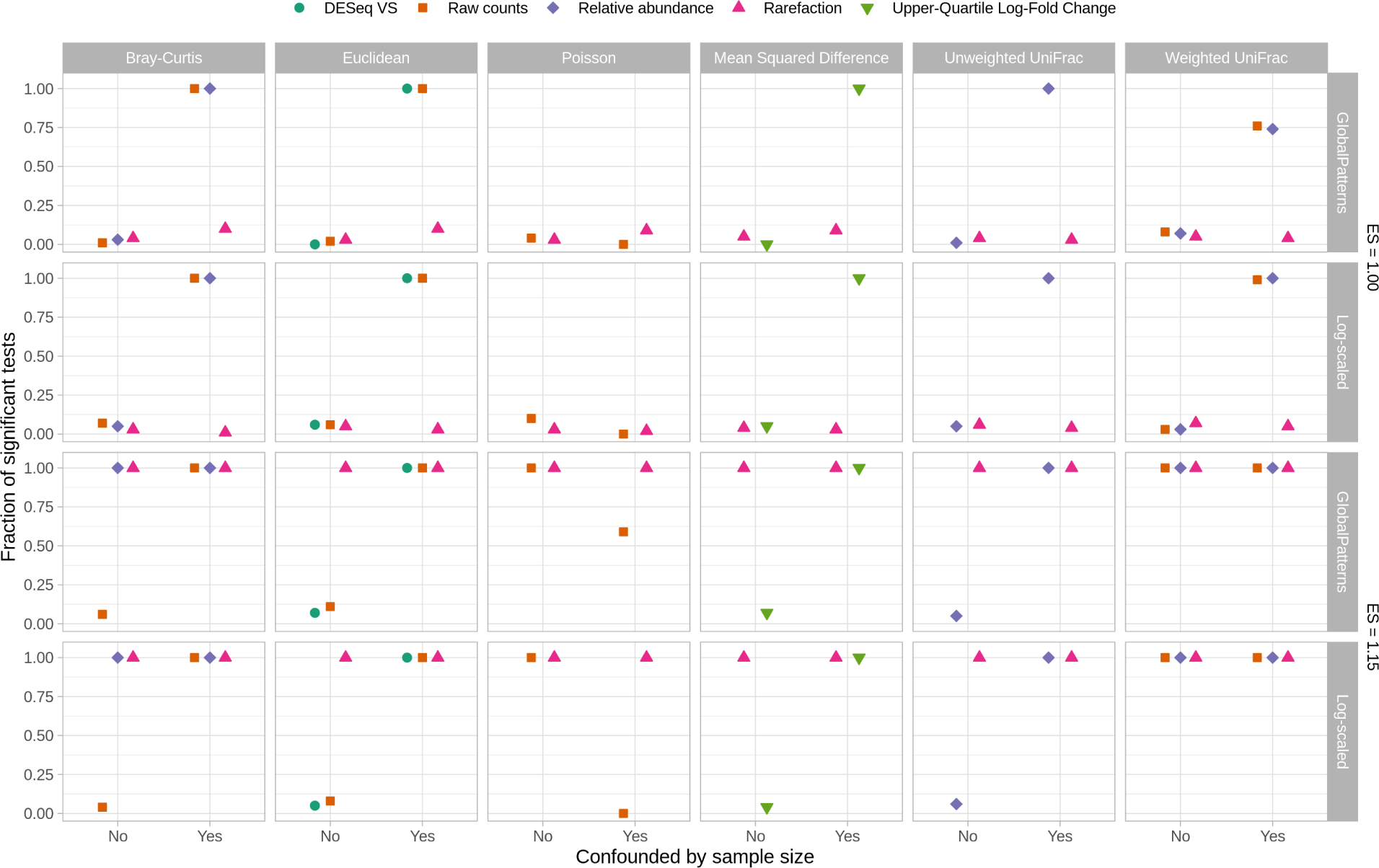
Rarefaction was consistently as good or better than all other normalization methods at controlling for Type I error and maximizing power to detect differences in treatment group using adonis2 regardless of whether sequencing depth was confounded by treatment group. Type I errors were assessed as the fraction of 100 simulations that yielded a significant P-value (i.e., less than or equal to 0.05) at an effect size of 1.00. Power was assessed as the fraction of 100 simulations that yielded a significant P-Value at an effect size of 1.15. Data are shown for a median sequencing depth (Ñ_L_) of 10,000 sequences when individual sequencing depths were sampled with replacement from the GlobalPatterns dataset or without replacement from the log-scaled distribution.

### 11. There was no consideration of effects of normalization methods on alpha diversity metrics, which is the traditional application of rarefaction

Rarefaction was originally proposed as a method for controlling uneven sequencing effort when comparing community richness values (5, 6). Thus it was surprising that WNWN did not consider the effect of the proposed normalizations on alpha-diversity metrics such as richness or Shannon diversity. Therefore, for each of the normalizations, I compared the richness and diversity of the two treatment groups. The DESeq Variance Stabilization normalized data were not included because the normalization produced negative values which were not compatible with calculations of richness or Shannon diversity. Also, data from the Upper-Quartile Log-Fold Change normalization were not used for richness calculations since the normalization returned the same richness values for each sample regardless of the treatment group. I assessed whether the alpha diversity was significantly different between the treatment groups for each iteration using the non-parametric Wilcoxon two-sampled test. I compared the risk of committing Type I errors and the power to detect differences by the different normalizations (Figures 16 and 17). For these analyses, I used the GlobalPatterns data with the random and confounded distribution of sequencing depths. Similar to the results in points 9 and 10, with the exception of rarefaction, the simulations using a confounded sequence depth distribution resulted in all of the replicates having a significant test. The power to detect differences in richness and diversity at effect sizes of 1.15 and greater with rarefaction was at least as high as any of the other normalizations.

**Figure 16.**
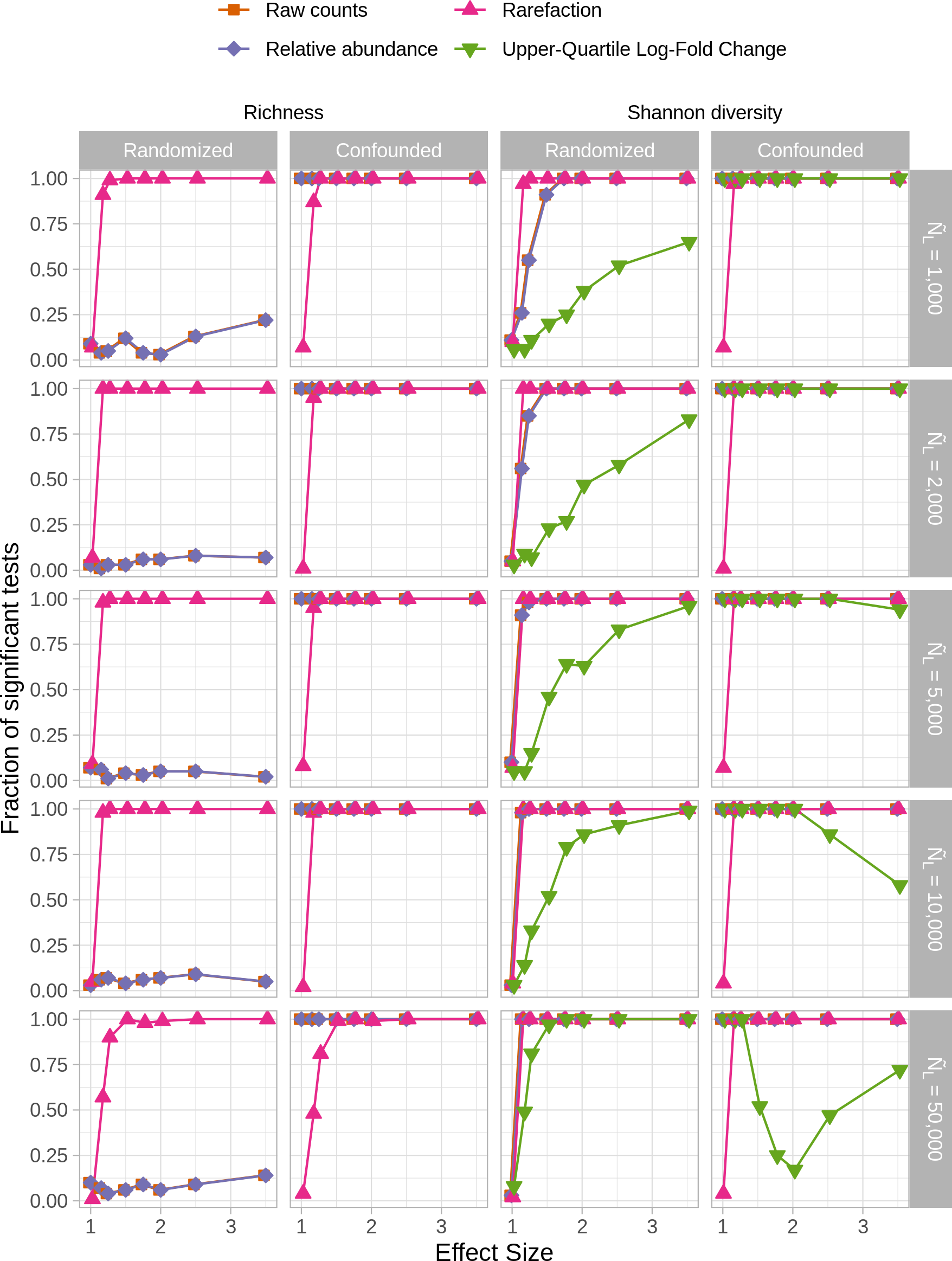
Rarefaction was consistently as good or better than all other normalization methods at controlling for Type I error and maximizing power to detect differences in treatment group using alpha-diversity metrics regardless of whether sequencing depth was confounded by treatment group. Statistical comparisons of OTU richness and Shannon diversity were performed using the non-parametric Wilcoxon two-sampled test. Type I errors were assessed as the fraction of 100 simulations that yielded a significant P-value (i.e., less than or equal to 0.05) at an effect size of 1.00. Power was assessed as the fraction of 100 simulations that yielded a significant P-Value at an effect size of 1.15. Data are shown for when the case when individual sequencing depths were sampled with replacement from the GlobalPatterns dataset.

**Figure 17.**
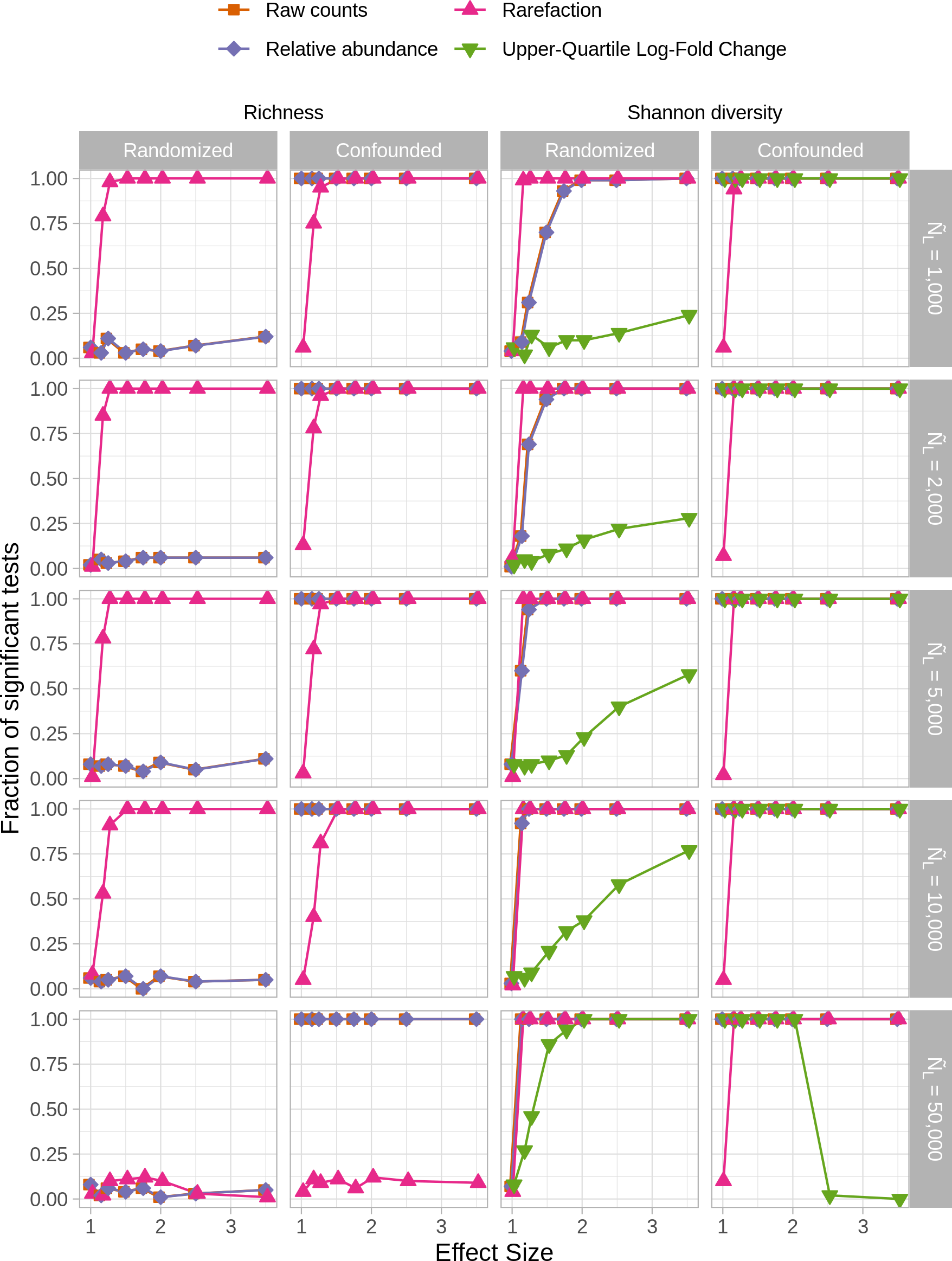
Rarefaction was consistently as good or better than all other normalization methods at controlling for Type I error and maximizing power to detect differences in treatment group using alpha-diversity metrics regardless of whether sequencing depth was confounded by treatment group. Statistical comparisons of OTU richness and Shannon diversity were performed using the non-parametric Wilcoxon two-sampled test. Type I errors were assessed as the fraction of 100 simulations that yielded a significant P-value (i.e., less than or equal to 0.05) at an effect size of 1.00. Power was assessed as the fraction of 100 simulations that yielded a significant P-Value at an effect size of 1.15. Data are shown for when the case when individual sequencing depths were sampled without replacement from the log-scaled distribution.

One odd result from this analysis was that the power to detect differences in richness at small effect sizes increased between 1,000 to 2,000 sequences with values of 0.91 and 1.00, respectively. The power then decreased with increasing sequencing effort to 0.57 with 50,000 sequences. This appeared to be because although the parent distributions had very different shapes, they shared a large number of rare taxa. When using rarefaction to compare the distributions at the size of the smallest distribution (i.e., 3,598,077 sequences), the feces parent distribution had and average of 1,559.65 OTUs and the ocean had an average of 1,335.00 OTUs; they shared an average of 894.66 OTUs. However, when comparing the distributions at 1,000 and 50,000 sequences, the Ocean distribution had greater richness than the feces distribution by 43.76 and 6.05, respectively. Therefore, although more replicates at lower effect sizes yielded a significant p-value at shallow rather than deeper sequencing depths for richness, the direction of the difference in richness was incorrect. This result underscores the challenges of using metrics based on presence-absence data like richness and Jaccard and Unweighted UniFrac distances to compare microbial communities.

## Discussion

The conclusions from WNWN have had a lasting impact on how researchers analyze microbiome sequence data. As I have demonstrated using WNWN’s simulation framework, their claims are not supported. The most important points that lead to the difference in our conclusions include the choice of clustering algorithm and arbitrarily selecting a sequencing effort threshold that was used to remove samples. Furthermore, the decision to evaluate the effectiveness of normalization methods based on clustering samples adds a layer of analysis that has been controversial and not widely used. Ultimately, the authors choice of the word “rarefying” has sewn confusion in the field because it is often used in place of the word “rarefaction”. As I have demonstrated there were numerous choices throughout the original study that resulted in rarefying/subsampling looking worse than it truly is. In fact, when the data of the current study are taken as a whole, rarefaction is the preferred approach. Short of obtaining the exact same number of sequences from each sample, rarefaction remains the best approach to control for uneven sequencing effort when analyzing alpha and beta diversity metrics. Although beyond the scope of the current study, the same principles would also support using rarefaction to control for uneven sequencing effort when calculating alpha and beta diversity metrics in shotgun metagenomic sequencing analyses.

Beyond the discussion of whether rarefaction is appropriate for analyzing microbiome data it is worth commenting on WNWN’s advice to use DESeq’s Variance Stabilization or edgeR’s Upper-Quartile Log-fold Change normalization strategies. These methods have been adopted from gene expression analysis to microbiome analysis. Gene expression analysis implicitly assumes that all samples have the same genes. While this might work in comparing healthy and diseased tissues from a cohort of patients, it does not generalize to those patients’ microbiota. Microbial populations are highly patchy in their distribution. Thus, a zero count for gene expression is more likely to represent a gene below the limit of detection whereas a zero count for a microbiome analysis is more likely to represent the true absence of the OTU. In fact, zeroes are less common in gene expression analyses. But because microbiome studies have so many zeroes, it is necessary to add a pseudo-count to all OTUs for both the edgeR and DESeq-based normalizations (L443 and L487, respectively). In WNWN, a pseudo-count of 1 was used. However, this value is arbitrary and the results can vary based on the choice of pseudo-count and the patchiness of the communities being analyzed. Since WNWN was published, compositional approaches have been proposed to account for uneven sequencing and to provide improved interpretability (14, 16–18, 46, 47). However, these methods also often require the use of pseudo-counts and are not actually insensitive to uneven sequencing (40). Rarefaction is preferred to these alternatives.

The choice of distance metric is a complicated question and the use of six different metrics in WNWN illustrates the challenges of selecting one. Within the ecology literature, Euclidean distances are widely avoided because joint absence is weighted the same as joint presence of taxa (31). As discussed in point 11, distances that are based on community membership (i.e., the presence or absence of taxa) performed worse than those than those that were based on community structure (i.e., their relative abundance). For this reason, the Unweighted UniFrac and other distances like the Jaccard or Sorenson distance coefficients should likely be avoided. The Poisson distance metric is largely novel to the microbial ecology literature and performed no better than the more traditional distances. In the current analysis, the phylogenetic Weighted UniFrac distance performed comparably to Bray-Cutis distances for clustering or differentiating between communities with adonis2. Regardless of the distance calculation employed, they all performed best when using rarefaction relative to the other normalization methods.

Although the authors claim that “Rarefying counts requires an arbitrary selection of a library size minimum that affects downstream inference” (page 8, column 1, point 3), in actual microbiome studies the selection of a sequencing depth is not as arbitrary as the authors claim. Rather, to avoid p-hacking, researchers pick a set of criteria where they will include or exclude samples prior to testing their data. Examples of criteria might include the presence of a large gap in the sequencing effort distribution, a desire to include poorly sequenced controls, or the *a priori* stipulation of a minimum sequencing effort. To mitigate concerns of arbitrary or engineered minimum library sizes, researchers should indicate the rationale for the threshold they selected.

WNWN performed a second set of simulations to address the effect of normalization method on the ability to correctly detect differential abundance of OTUs that were randomly selected to have their relative abundances changed (i.e., “Simulation B”). Re-addressing this set of simulations is beyond the scope of the current analysis and others have already contributed critiques (25). However, many of the same concerns addressed above would apply. Perhaps most important is the fact that if the relative abundance of several OTUs increase, then the relative abundance of all other OTUs would necessarily decrease because the data are compositional (47). Thus, one would expect every OTU to be differentially abundant. Because of this, it is not possible to truly modify the abundance of a set of OTUs independent of all other OTUs. This is an important limitation of tests of methods attempting to detect differentially abundant OTUs. Regardless, a form of rarefaction could still be employed for detecting differential abundance. One could subsample the data, perform the statistical test, and identify the differentially abundant OTUs. This process could be repeated and the results synthesized. In my experience the largest differences between subsamples are for low relative abundances OTUs, which are unlikely to be statistically significant.

In a parallel set of analyses I have used alternative simulation and evaluation strategies to look more closely at rarefaction and its alternatives (40) and the practice of filtering low abundance sequences (4). The results of those studies are similar to this analysis. These studies demonstrate that it is actually sequence and OTU filtering that “requires the omission of available valid data” and that rarefaction is the only available method for controlling the effects of uneven sequencing. Far from being inadmissible, rarefaction of unfiltered datasets yields the most robust results.

## Methods

### Code availability

A git repository containing all of the code needed to reproduce this study is available at https://www.github.com/SchlossLab/Schloss_WNWN_XXXXX_2023. All simulations and analyses were performed using R and bash scripts using Snakemake (v7.24.0) to track dependencies and automate the pipeline and Conda (v4.12.0) and mamba (v1.1.0) to specify software and package versions (see workflow/envs/nr-base.yml in the repository). This manuscript was written as an R markdown document and rendered using the rmarkdown R package (v2.18). All figures were generated using dplyr (v1.1.0) and ggplot2 (v3.4.2) from the tidyverse metapackage (v1.3.2) and ggtext (v0.1.2) and ggh4x (v0.2.4) within R (v.4.2.2; see workflow/envs/nr-modern.yml in the repository).

### Reproducing WNWN Protocol S1

The compressed directory published with WNWN as Protocol S1 is available on the *PLOS Computational Biology* website with WNWN at https://doi.org/10.1371/journal.pcbi.1003531.s001. To render the R markdown files to HTML files using software and packages as similar as possible to that of WNWN, I created an environment (see workflow/envs/nr-s1.yml in this study’s repository) that contained versions as close to those indicated in the pre-rendered files. The most important packages for the reproduction of the cluster analysis included DESeq (my version: v1.39.0 vs. WNWN version: v1.14.0), edgeR (v3.30.0 vs. v3.4.0), cluster (v2.1.0 vs v1.14.4), phyloseq (v1.32.0 vs. v1.6.0), and PoiClaClu (v1.0.2.1 vs. 1.0.2). In addition, I used R v4.0.2 whereas the WNWN analysis used v3.0.2. Finally, to render the simulation-cluster-accuracy-server.Rmd file to HTML, it was necessary to apply a patch to the code. This patch set the path to the location where the figures should be saved and removed the R code that deleted all objects in the environment. Both changes were necessary to better organize the project and had no substantial bearing on the content of the rendered HTML file.

### Simulated communities

The code contained within simulation-cluster-accuracy-server.Rmd was spilt into individual R scripts and modified to make the generation of the simulated communities more modular and scalable. The code for generating the simulated communities was run using the nr-s1 Conda environment. The ocean and feces parent distributions were generated as described above. Five variables were altered to simulate each set of communities. First, eight mixing fractions were used to manipulate the effect size between the two simulated treatment groups with 40 samples per group. These were the same effect sizes used in the WNWN study (1.00 [i.e., no difference], 1.15, 1.25, 1.50, 1.75, 2.00, 2.50, and 3.50). Second, the number of sequences in each of the 80 samples was either randomly selected from the 26 library sizes contained within the GlobalPatterns dataset or from a log distribution of 80 sequencing depths evenly distributed between the most shallow (58,688 sequences) and deeply (2,357,181 sequences) sequenced samples in the GlobalPatterns dataset (Figure 10). Third, the resulting 80 sequencing depths were scaled so that the median sequencing depth across the 80 samples was either 1,000, 2,000, 5,000, 10,000, or 50,000 sequences. Fourth, the sequencing depth of each sample was either randomly assigned to each treatment group or assigned so that those depths less than or greater than the median were assigned to separate treatment groups. Finally, each set of conditions was replicated 100 times. These five parameters resulted in 1,600 simulated datasets (8 effect sizes x 2 sequencing depth models x 5 sequencing depths x 2 sequencing depth assignment models x 100 replicates).

### Generation of distances between communities

Next, each simulation was further processed by filtering rare OTUs, to normalize uneven sequencing depths, and to calculate ecological distances between samples. Again, the code contained within simulation-cluster-accuracy-server.Rmd was spilt into individual R scripts and modified to make the generation of the simulated communities more modular and scalable within the nr-s1 Conda environment. First, each simulation was either filtered to remove rare OTUs or left unfiltered. As was done in the WNWN study, filtering was performed in two steps: (1) any OTUs whose total abundance was less than 3 across all 80 samples and that did not appear in at least 3 samples and (2) any OTU that did not have more than 1 sequence in more than 5% of the 80 samples (i.e., 4 samples) and that did not have a total abundance across the 80 samples greater than one half of the number of samples in each community type (i.e., 20) were removed. Second, OTU data was normalized by one of six approaches: (1) non-normalized raw counts, (2) relative abundance, (3) DESeq Variance Stabilization, (4) Upper-Quartile Log-Fold Change normalization using edgeR, (5) a single subsampling to a common number of sequences, (6) rarefaction with 100 randomizations to a common number of sequences. Although the data were not presented, the WNWN code and my code also normalized the counts using trimmed mean of M-values (TMM) and relative log expression (RLE) in edgeR. The level of subsampling and rarefaction was selected based on the size of the sample whose sequencing depth was at the 0 (i.e., included all of the samples), 5, 10, 15, 20, 25, and 40th percentiles. Finally, the normalized simulated datasets were used to calculate pairwise distances between the 80 samples using seven different calculations: Bray-Curtis, Euclidean, Poisson, Mean Squared Difference, Unweighted UniFrac, Weighted Unifrac, and Biological Coeffcient of Variation (BCV); however, data using BCV were not discussed in this or the WNWN analysis. For the rarefaction normalized data, each of the 100 subsampled datasets were used to calculate a distance matrix. The mean of the 100 distance matrices was used as the distance matrix for the rarefaction data. As indicated in the Results section, there were combinations of normalization and distance methods that were not compatible (e.g., DESeq Variance Stabilization normalization with Bray-Curtis distances). In such cases, distance matrices were coded as NA. My choice of combinations of normalizations and distances to visualize was determined by the WNWN analysis except where noted. For each simulated dataset there were 280 possible combinations of filtering, normalization, and distance methods (2 filtering methods x 20 normalization methods x 7 distance methods).

### Analysis of distances between communities

Two general strategies were used to analyze the pairwise distances in each distance matrix. First, by recycling the the code contained within simulation-cluster-accuracy-server.Rmd in the nr-s1 Conda environment, samples were assigned to one of two clusters using partitioning around the medoid (PAM) with the pam function from the cluster R package, K-means clustering with the kmeans function from the stats base R package, and hierarchical clustering with the hclust and cutree functions from the stats base R package. The accuracy of the clustering was measured as the fraction of the 80 samples assigned to the correct cluster. As in WNWN, if samples were removed because the number of sequences in them fell below the threshold for subsampling or rarefaction, then the accuracy could be below 50%. For the effect size of 1.00, all of the samples should have been assigned to one group; however, because they were forced into two groups there would be 41 “correctly” assigned samples of the 80 samples (i.e., an accuracy of 0.51). Second, the distance matrices were analyzed for significant different centroids using the adonis2 function from the vegan package (v2.6-4) in the nr-modern Conda environment. The fraction of significant tests was the fraction of the 100 replicate simulations that had a p-value less than or equal to 0.05. The Type I error rate for a condition was the fraction of significant tests when the effect size was 1.00. The power was the fraction of significant tests at other effect sizes.

### Analysis of richness and diversity between communities

Using the nr-s1 Conda environment, I measured the richness and Shannon diversity using the normalized OTU counts. Richness was measured by counting the number of OTUs for each sample across the simulations and the Shannon diversity using their relative abundance using the commonly used formula (31). For the rarefaction data, each of the 80 samples in the 100 subsampled datasets were used to calculate the richness and diversity. The mean across the 100 subsamplings was used as the value for the rarefaction data across each of the 80 samples. To assess differences between the two treatment groups, the richness and diversity values were compared using the two-sample Wilcoxon non-parametric test with the wilcox.test from the stats base R package. Similar to the analysis using adonis2, the fraction of significant tests was the fraction of the 100 replicate simulations that had a p-value less than or equal to 0.05. The Type I error rate for a condition was the fraction of significant tests when the effect size was 1.00. The power was the fraction of significant tests at other effect sizes.

## Acknowledgements

I am grateful to Dr. Paul McMurdie and Dr. Susan Holmes for publishing the source code that they used to conduct their analysis as R markdown files in the supplement to WNWN. It would not have been possible for me to replicate and build upon their simulations without their code.

## Supplementary text

Quotes from selected papers that cited WNWN and conflated “rarefying”/“rarefy” and “rarefaction”. Many of these papers also attributed “rarefaction” to WNWN. Rarefying, rarefy, and rarefaction are bolded to make the words easier to identify. All references have been updated to reflect the numbering in the current study. Citations to WNWN are indicated by (24). Quotes are ordered chronologically.

- “Additionally, **rarefaction** has recently been shown to introduce errors in analyses, and alternatives to rarefaction have been proposed (24)” - Quoted from (48). Implies that rarefaction and WNWN’s rarefying are the same thing.
- “**Rarefaction** is analytically problematic and poses multiple statistical problems: (i) omission of available valid data, (ii) the estimation of overdispersion is more difficult due to data loss, (iii) loss of power (type II error), (iv) dependence on an arbitrary threshold and (v) additional uncertainty due to the randomness in **rarefaction** (24).” - Quoted from (49). Implies that rarefaction and WNWN’s rarefying are the same thing.
- “McMurdie and Holmes (24) penalized the **rarefying** technique for dropping the lowest fifteenth percentile of sample library sizes in their simulations by counting the dropped samples as”incorrectly clustered.” Because the 15th percentile was used to set **rarefaction** depth, this capped clustering accuracy at 85%.” - Quoted from (25). An example of a paper from the QIIME development community using rarefaction and rarefying interchangeably.
- “**Rarefaction** has a limited ability in this regard since the total sum constraint still exists after **rarefaction**. In addition, it suffers from a great power loss due to the discard of a large number of reads (24).” - Quoted from (50). Implies that rarefaction and WNWN’s rarefying are the same thing.
- “Another popular normalization approach is **rarefaction**, which consists on subsampling the same number of reads for each sample so that all samples have the same number of total counts. **Rarefaction** is not recommended because it entails the loss of important information (24).” - Quoted from (51). Implies that rarefaction and WNWN’s rarefying are the same thing.
- “Unfortunately, **rarefaction** is neither justifiable nor necessary, a view framed statistically by McMurdie and Holmes (24) in the context of comparison of relative abundances.” - Quoted from (21). Implies that rarefaction and WNWN’s rarefying are the same thing.
- “Some studies perform **rarefaction** to adjust for differences in library size due to unexhaustive metagenomic sampling. Although several pipelines provide this functionality, it has been found inadmissible for metagenomics microbiome studies as it discards many reads leading to decreased sensitivity in differential abundance testing (24) and biased estimates for alpha diversity (21).” - Quoted from (52). Implies that rarefaction and WNWN’s rarefying are the same thing.
- “Another widespread practice is to **rarefy** the count data to force the samples to have the same number of total sequence reads (13), at the expense of discarding vast amounts of information (24).” - Quoted from (53). Hughes and Hellmann (13) describe traditional rarefaction with multiple subsamplings.
- “Note that the terms **rarefying** and **rarefaction** are used interchangeably in microbiome literature (24). **Rarefying** was first recommended for microbiome data to deal with rare taxa (54), which impact some measures of alpha and beta diversities (24).” - Quoted from (14). As the quote indicates they use rarefying and rarefaction interchangeably although the cited WNWN indicates that they are not the same. In addition, the first review article to advocate the use of rarefaction was Hughes et al. (12) and examples of its use can be found earlier (55, 56);
- “Ways to tackle this include total library size normalization and **rarefaction**, with both remaining debated to date (24, 25).” - Quoted from (57). Implies that rarefaction and WNWN’s rarefying are the same thing.
- “**Rarefaction** is a widely used normalization technique that involves the random subsampling of sequences from the initial sample library to a selected normalized library size. This process is often dismissed as statistically invalid because subsampling effectively discards a portion of the observed sequences, yet it remains prevalent in practice and the suitability of **rarefying**, relative to many other normalization approaches, for diversity analysis has been argued” - Quoted by (30). Uses rarefaction and rarefying interchangeably. This paper describes a new method as “repeated rarefying”, which is the traditional rarefaction approach.
- “Confronted with technical variation as well as the overall increase in raw sequencing data generated per sample over the years, **rarefaction** (or downsampling) was suggested to standardize within and across dataset comparisons . . . However, sequencing depth-based downsizing procedures were soon criticized, not only for being wasteful and discarding information on low-abundant taxa (24), but also for being unsuited when applied to communities characterized by substantial variation in cell density (58).” - Quoted by (59). Implies that rarefaction and WNWN’s rarefying are the same thing.
- “Some have suggested that data should be **rarefied** (23, 25) to a common sampling depth, typically to the level of the sample with fewest sequences, while others argue that such **rarefaction** is ‘inadmissible’ and favor approaches that transform or scale sequence counts (24, 47).” - Quoted by (60). Uses rarefied and rarefaction interchangeably and implies that both are the same as WNWN’s rarefying.
- “**Rarefaction** has been criticized for wasting data since we effectively remove a portion of the data in the downsampling procedure (24).” - Quoted by (61). Here and elsewhere in this paper the authors use rarefaction and rarefying interchangeably and implies that both are the same as WNWN’s rarefying.
- “**Rarefying** (also referred to as **rarefaction**) is a popular but widely criticized technique for correcting uneven sequencing depths. It involves randomly discarding counts in samples until all samples have the same predefined number of total counts.” - Quoted by (62). As the quote indicates they use rarefying and rarefaction interchangeably although the cited WNWN indicates that they are not the same.
- “One commonly used method is to **rarefy** the data; that is, ASVs or OTUs within a sample are randomly subsampled without replacement to a preselected depth that is the same across all samples. The outcome of this is that all samples will have the same number of ASVs and any samples with fewer ASVs than the **rarefaction** level will be removed from the dataset. The level for **rarefaction** can be decided using a **rarefaction** curve, a method in which each sample is subsampled at multiple levels (e.g. 1,000 reads, 2,000 reads, 3,000 reads. . .), and the number of unique features or another metric of individual sample diversity of each sample at each level is measured and plotted. When the plot begins to level off after an initial climb up, the corresponding number of sequences indicates an appropriate sampling depth. The appropriate number to **rarefy** must then be balanced with the number of samples that may be dropped from the dataset which do not meet that minimum. An advantage of **rarefaction** is that it may be a more appropriate measure of very low-abundance (”rare”) ASVs. This can in turn increase the accuracy of the data, as low biomass samples often have contamination and quality concerns (63). There are also disadvantages to this method, the most obvious of which is the discarding of valuable data. Clearly, this is less than ideal as the researcher must pay for the samples and sequences, and in cases where the samples are very valuable or difficult to obtain the loss of data may be destructive to the overall experimental integrity. Additionally, the loss of statistical power by removing sequences from a sample could lead to a loss of differences between two samples (24). The statistical consequences extend beyond this, as **rarefying** equalizes sample variance by adding artificial uncertainty (24).” - Quoted by (64). The authors use rarefaction and rarefying interchangeably and implies that both are the same as WNWN’s rarefying.

